# Ribosomal quality control factors inhibit repeat-associated non-AUG translation from GC-rich repeats

**DOI:** 10.1101/2023.06.07.544135

**Authors:** Yi-Ju Tseng, Indranil Malik, Xiexiong Deng, Amy Krans, Karen Jansen-West, Elizabeth M.H. Tank, Nicolas B. Gomez, Roger Sher, Leonard Petrucelli, Sami J. Barmada, Peter K. Todd

**Author notes:** To whom correspondence should be addressed: Peter K. Todd, MD, PhD, Department of Neurology University of Michigan 4005 BSRB, 109 Zina Pitcher Place Ann Arbor, MI 48109.

## Abstract

A GGGGCC (G4C2) hexanucleotide repeat expansion in *C9ORF72* causes amyotrophic lateral sclerosis and frontotemporal dementia (C9ALS/FTD), while a CGG trinucleotide repeat expansion in *FMR1* leads to the neurodegenerative disorder Fragile X-associated tremor/ataxia syndrome (FXTAS). These GC-rich repeats form RNA secondary structures that support repeat-associated non-AUG (RAN) translation of toxic proteins that contribute to disease pathogenesis. Here we assessed whether these same repeats might trigger stalling and interfere with translational elongation. We find that depletion of ribosome-associated quality control (RQC) factors NEMF, LTN1, and ANKZF1 markedly boost RAN translation product accumulation from both G4C2 and CGG repeats while overexpression of these factors reduces RAN production in both reporter cell lines and C9ALS/FTD patient iPSC-derived neurons. We also detected partially made products from both G4C2 and CGG repeats whose abundance increased with RQC factor depletion. Repeat RNA sequence, rather than amino acid content, is central to the impact of RQC factor depletion on RAN translation - suggesting a role for RNA secondary structure in these processes. Together, these findings suggest that ribosomal stalling and RQC pathway activation during RAN translation elongation inhibits the generation of toxic RAN products. We propose augmenting RQC activity as a therapeutic strategy in GC-rich repeat expansion disorders.

**Graphical Abstract:** 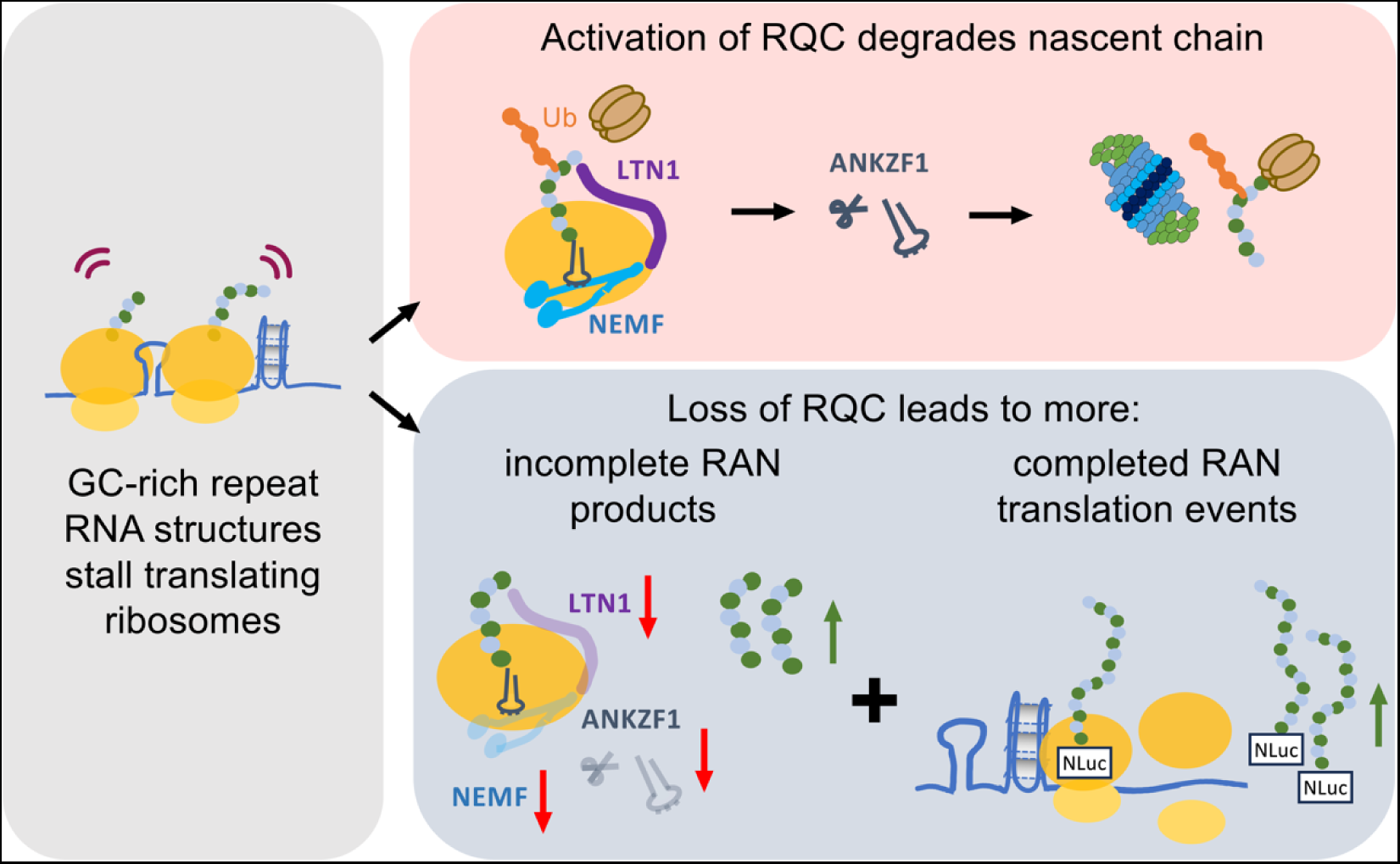

## INTRODUCTION

Short Tandem Repeat (STR) expansions cause more than 50 human neurodegenerative, neurodevelopmental, and neuromuscular disorders (1–6). Several STR expansions trigger a process known as repeat-associated non-AUG (RAN) translation, which is a non-canonical initiation process whereby proteins are generated from repeats without an AUG start codon (7–14). RAN translation-generated proteins accumulate within neurons and in patient tissues and elicit toxicity in both cellular and animal disease models (9, 15–20). As at least 10 different neurodegenerative disease-associated STRs support RAN translation, there is significant interest in identifying selective inhibitors and regulators of this non-canonical translation process (6, 13, 20–30). However, to date, the precise mechanisms that regulate RAN translation remain largely unknown.

Amyotrophic lateral sclerosis (ALS) and frontotemporal dementia (FTD) are inexorably progressive and fatal neurodegenerative disorders. The most common genetic cause of FTD and ALS is an expanded GGGGCC (G4C2) hexanucleotide repeat within the first intron of *C9ORF72* (C9FTD.ALS)(31–33). Fragile X-associated tremor/ataxia syndrome (FXTAS) is similarly a progressive adult-onset neurodegenerative disorder without effective treatments. FXTAS results from a transcribed CGG repeat expansion in the 5’ UTR of *FMR1* that affects about 1/5000 people worldwide (34–37). The repeats that cause both C9ALS/FTD and FXTAS are highly GC-rich and are thought to form strong secondary structures as RNA, including RNA hairpins and G quadruplexes (38–43). Moreover, both repeat sequences support RAN translation (14, 17, 20, 25, 44–47). Without an AUG start codon, RAN translation can occur in all possible reading frames from both sense and antisense transcripts (46, 48). For example, G4C2 repeats in *C9ORF72* generate GA (glycine-alanine, GA; +0 frame), GP (glycine-proline, GP; +1 frame), and GR (glycine arginine, GR; +2 frame) dipeptide repeat proteins (DPRs) while CGG repeats in *FMR1* generate polyG (glycine; +1 frame), polyA (alanine; +2 frame), and polyR (arginine; +0 frame) homopolymeric peptides from the sense transcripts (46, 49). However, the efficiency of RAN translation differs significantly across reading frames for each repeat, such that GA DPRs are produced more readily than GP or GR from G4C2 repeats, and polyG and polyA are more efficiently generated than polyR peptides from CGG repeats (17, 20, 44, 45, 49). While initial studies suggested that these differences might be due solely to near-cognate initiation sites in the sequences surrounding the repeats, more recent work suggests that these differences in translational efficiency persist even when each product is generated from an AUG codon in an ideal Kozak context (44, 49–52). This suggests that differences in translational elongation efficiency may be critically important to RAN translation efficiency and as such could represent a therapeutic target.

Ribosomes stall during translational elongation in response to many cues, including RNA damage, mRNA secondary structures, the charge of the nascent polypeptide chain, and cellular stress (53–58). Stalled ribosomes trigger a series of mRNA and protein surveillance pathways known collectively as ribosome-associated protein quality control (RQC) (59–65). These quality control pathways ensure the fidelity of protein synthesis and degrade incompletely generated polypeptides and the mRNAs from which they are derived. Stalled ribosomes are sensed by RING-domain E3 ubiquitin ligase, zinc finger protein 598 (ZNF598, yeast Hel2), and receptor for activated C kinase 1 (RACK1, yeast Asc1) to dissociate the 80S ribosome (66–68). Alternatively, stalling at the 3’ end of mRNA is detected by the mRNA surveillance and ribosome rescue factor Pelota (PELO, yeast Dom34) and HBS1-like translational GTPase (HBS1L, yeast Hbs1) (69–71). The PELO:HBS1L complex subsequently recruits ATP binding cassette subfamily E member 1 (ABCE1, yeast Rli1) to disassemble the ribosome (72, 73). Separation of 80S subunits releases 40S and 60S subunits. The 40S subunits contain the truncated mRNA, which is degraded by 5’ - 3’ exoribonuclease (XRN1) and the exosome complex (74). 60S subunits entrapped with tRNA-bound nascent polypeptide leads are engaged by the ribosomal quality control (RQC) complex, which targets partially made nascent polypeptides for proteasomal degradation (62, 64, 75–79). Assembly of the RQC complex is initiated by the binding of nuclear export mediated factor (NEMF, yeast Rqc2), which recruits and stabilizes binding by the E3 ligase listerin (LTN1, yeast Ltn1) (79–83). NEMF also synthesizes carboxy-terminal alanine and threonine tails (CAT-tails) on partially made polypeptide chains (84–87). This pushes amino acids on the nascent chain out from the ribosomal exit tunnel and triggers their ubiquitination by LTN1 (78, 88–91). The ubiquitin chains signal for recruitment of the valosin-containing protein (VCP, yeast Cdc48) (92, 93). The nascent polypeptide chain is then released by ankyrin repeat and zinc finger peptidyl tRNA hydrolase 1 (ANKZF1, yeast Vms1) and VCP targets the ubiquitylated polypeptide chain for proteasomal degradation (94–96).

Previous studies demonstrated that ribosomes stall during the synthesis of C9 ALS/FTD-associated GR and PR dipeptide proteins (97–99). These stalling events were not seen on similarly sized but uncharged DPR repeats such as GA and were therefore ascribed to positively charged arginine residues within these nascent polypeptide chains. Further studies suggest that the GR protein may also interact with protein surveillance pathways through other means than its directed translation (98, 100, 101). However, these studies were not done in the context of the GC-rich RNA repeats or in the absence of AUG start codons as occurs in RAN translation. As such, whether the dynamics of translation and/or the potential for repeat RNA structures to elicit ribosomal stalling and RQC pathway engagement remains largely unexplored.

In this study, we performed a targeted RAN translation modifier screen at two different GC-rich short tandem repeats with factors involved in the mRNA and protein surveillance pathways. We find that NEMF, LTN1, and ANKZF1 from the RQC pathway all act as significant modifiers of RAN translation from GC-rich repeat RNAs. This is true for the generation of both GA and polyG polypeptides, even though both products contain no runs of charged amino acids. Consistent with translational stalling in these reading frames, we detected partially made N-terminal peptides containing GA from G4C2 repeats and polyG from CGG repeats, and depletion of these RQC factors enhance the accumulation of these truncated repeat peptides. We see a similar effect that the RQC factor impacts on the accumulation of RAN products in patient-derived neurons and find that altering NEMF directly impacts repeat-associated toxicity in a *Drosophila* model system. Taken together, our results suggest that RQC factor engagement during translational elongation through GC-rich short tandem repeats directly impacts both RAN translation efficiency and product generation through a repeat RNA structure-dependent mechanism. Moreover, as modulating RQC factor abundance modulates RAN product abundance, targeted augmentation of RQC activity is an intriguing target for further therapeutic evaluation.

## MATERIALS AND METHODS

### Plasmids

Plasmids and cloning primers used in this study and their sources are listed in **Table 1**. Primers and synthesized fragments used for cloning are listed in **Table 2**. ATG-V5-GA_69_-NLuc-3xFLAG was generated by the insertion of an ATG-V5 fragment flanked with NheI/EcoRV upstream of Intron-(GGGGCC)_70_-NLuc-3xFLAG (GA frame). ATG-V5-+1(CGG)_101_-NLuc-3xFLAG was generated by the insertion of an ATG-V5 sequence flanked by EcoRI and NarI upstream of +1(CGG)_100_-NLuc-3xFLAG (polyG frame). ATG-V5-(GGN)_103_ sequence was synthesized by GeneWiz with flanking restriction enzymes of EcoRI (5’) and XhoI (3’). The synthesized sequence was inserted into the ATG-V5-+1(CGG)_101_-NLuc-3xFLAG plasmid with EcoRI/XhoI to replace the ATG-V5-+1(CGG)_101_ sequence. pBI-dsRED/hNEMF R86S was a gift from Roger Sher. WT hNEMF was generated by mutating the serine at site 86 to an arginine using Q5 site-directed mutagenesis (NEB, E0554S). pBI-dsRED empty vector was cloned using annealing primers flanked by NheI and EcoRV. pCMV3TAG8li-hRNF160-3FLAG was a gift from Martin Dorf (Addgene plasmid #159138). pCMV6-hANKZF1-Myc-FLAG was purchased from Origene, RC201054. All other reporter sequences have been previously published (44, 49).

**Table 1.** Plasmids information

| Plasmid | Source | Cat# |
| --- | --- | --- |
| pcDNA3.1-ATG-NLuc-3xFLAG | Kearse et al., 2016 | N/A |
| pcDNA3.1-Intron-(GGGGCC)<sub>3</sub>-NLuc-3xFLAG (+0 polyGA frame) | Green et al., 2017 | N/A |
| pcDNA3.1-Intron-(GGGGCC)<sub>35</sub>-NLuc-3xFLAG (+0 polyGA frame) | Green et al., 2017 | N/A |
| pcDNA3.1-Intron-(GGGGCC)<sub>70</sub>-NLuc-3xFLAG (+0 polyGA frame) | Green et al., 2017 | N/A |
| pcDNA3.1-Intron-(GGGGCC)<sub>70</sub>-NLuc-3xFLAG (+1 polyGP frame) | Green et al., 2017 | N/A |
| pcDNA3.1-Intron-(GGGGCC)<sub>70</sub>-NLuc-3xFLAG (+2 polyGR frame) | Green et al., 2017 | N/A |
| pcDNA3.1-(CGG)<sub>100</sub>-NLuc-3xFLAG (+0 polyR frame) | Kearse et al., 2016 | N/A |
| pcDNA3.1-(CGG)<sub>100</sub>-NLuc-3xFLAG (+1 polyG frame) | Kearse et al., 2016 | N/A |
| pcDNA3.1-(CGG)<sub>100</sub>-NLuc-3xFLAG (+2 polyA frame) | Kearse et al., 2016 | N/A |
| pcDNA3.1-ATG-V5-NLuc-3xFLAG | Kearse et al., 2016 | N/A |
| pcDNA3.1-ATG-V5-(GGGGCC)<sub>69</sub>-NLuc-3xFLAG (polyGA frame) | This paper | N/A |
| pcDNA3.1-ATG-V5-(CGG)<sub>101</sub>-NLuc-3xFLAG (polyG frame) | This paper | N/A |
| pcDNA3.1-ATG-V5-(GGN)<sub>103</sub>-NLuc-3xFLAG (polyG peptides) | This paper | N/A |
| pBI-CMV2-dsRED | This paper | N/A |
| pBI-CMV4-hNEMF/CMV2-dsRED | This paper | N/A |
| pCMV-FLAG | Sigma | E7908 |
| pCMV-3TAG8li-hRNF160-3xFLAG (hLTN1) | Addgene | 159138 |
| pCMV6-hANKZF1-Myc-FLAG | Origene | RC201054 |

**Table 2.**
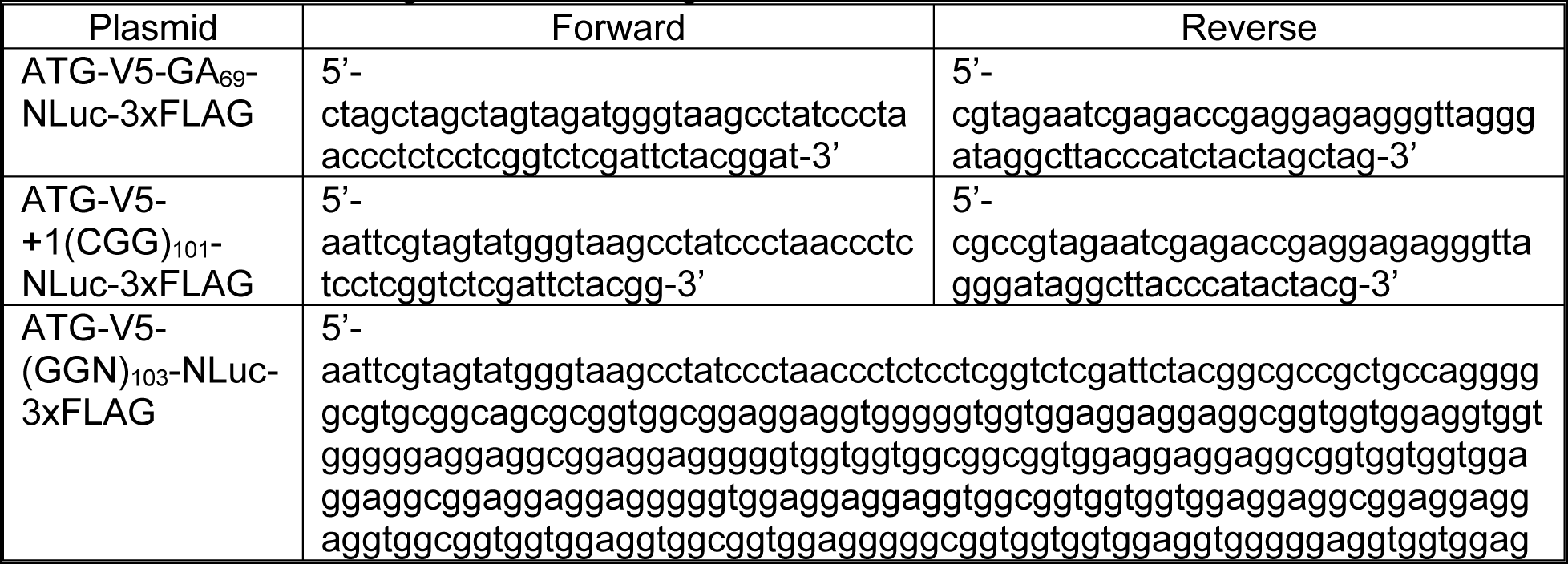
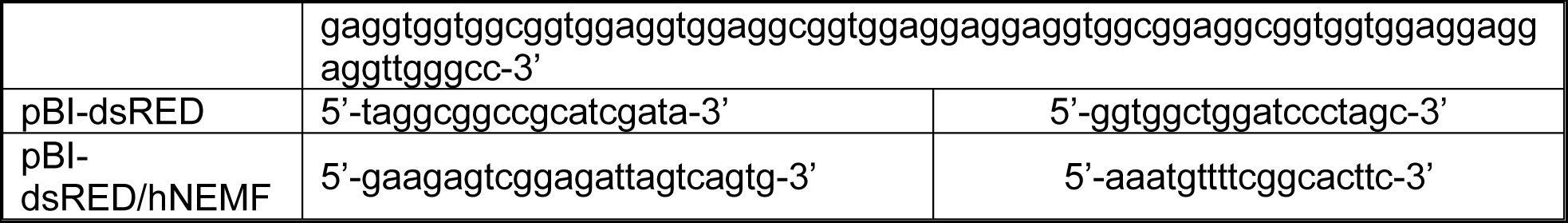
Primers and fragments for cloning

### RNA T7 synthesis

pcDNA3.1(+) plasmids containing nano-luciferase and 3xFLAG (NLuc-3xFLAG) reporters were linearized after the 3’ FLAG tag with PspOMI. The efficiency of restriction enzyme digestion was confirmed with a DNA agarose gel. Linearized DNA plasmids were cleaned and concentrated with DNA Clean & Concentrator-25 (Zymo Research, D4033). RNAs were *in vitro* transcribed with HiScribe T7 ARCA mRNA Kit with tailing (NEB, E2060S) following the manufacturer’s recommended protocol. mRNAs were then cleaned and concentrated with RNA Clean & Concentrator-25 Kit (Zymo Research, R1017) and run on a denaturing glyoxal RNA gel to verify mRNA size and integrity. Transcribed RNA sequences are shown in **Table 3**.

### Cell culture and transfections

HEK293 cells were maintained at 37°C, 5% CO_2_ in DMEM with high glucose (Gibco, 11965118) supplemented with 9% fetal bovine serum (50 mL FBS added to 500 mL DMEM; Bio-Techne, S11150). For siRNA transfection, HEK293 cells were seeded at 2x10^5^ cells/mL with 100 μL in 96-well or 500 μL in 24-well plates. Cells were then transfected with siRNA at 1 nM/well in a mixture with Lipofectamine RNAiMax (Thermo Fisher, 13778075) following the manufacturer’s recommended protocol on the same day while seeding the cells. siRNA used for this paper are listed in **Table 4**. For effector plasmid transfection, 24 hr after seeding the cells at 50-60% confluency, 100 ng and 500 ng of effectors were transfected in each well of 96-well and 24-well plate, respectively, with FuGeneHD (Promega, E2312), at a 3:1 ratio of FuGeneHD to DNA. Plasmid reporters were transfected into 70-80% confluent cells 48 hr post-seeding. Each well of a 96-well and 24-well plate was transfected with 50 ng and 500 ng NL reporter DNA, respectively, with FuGeneHD as described above. Cells were harvested 24 hr post-transfection for analysis. For mRNA transfection, T7 synthesized mRNA was transfected in cells at 70-80% confluency with TransIT-mRNA Transfection Kit (Mirus, MIR 2256) following the manufacturer’s recommended protocol. Cells were harvested 24 hr post-transfection for analysis.

**Table 3.**
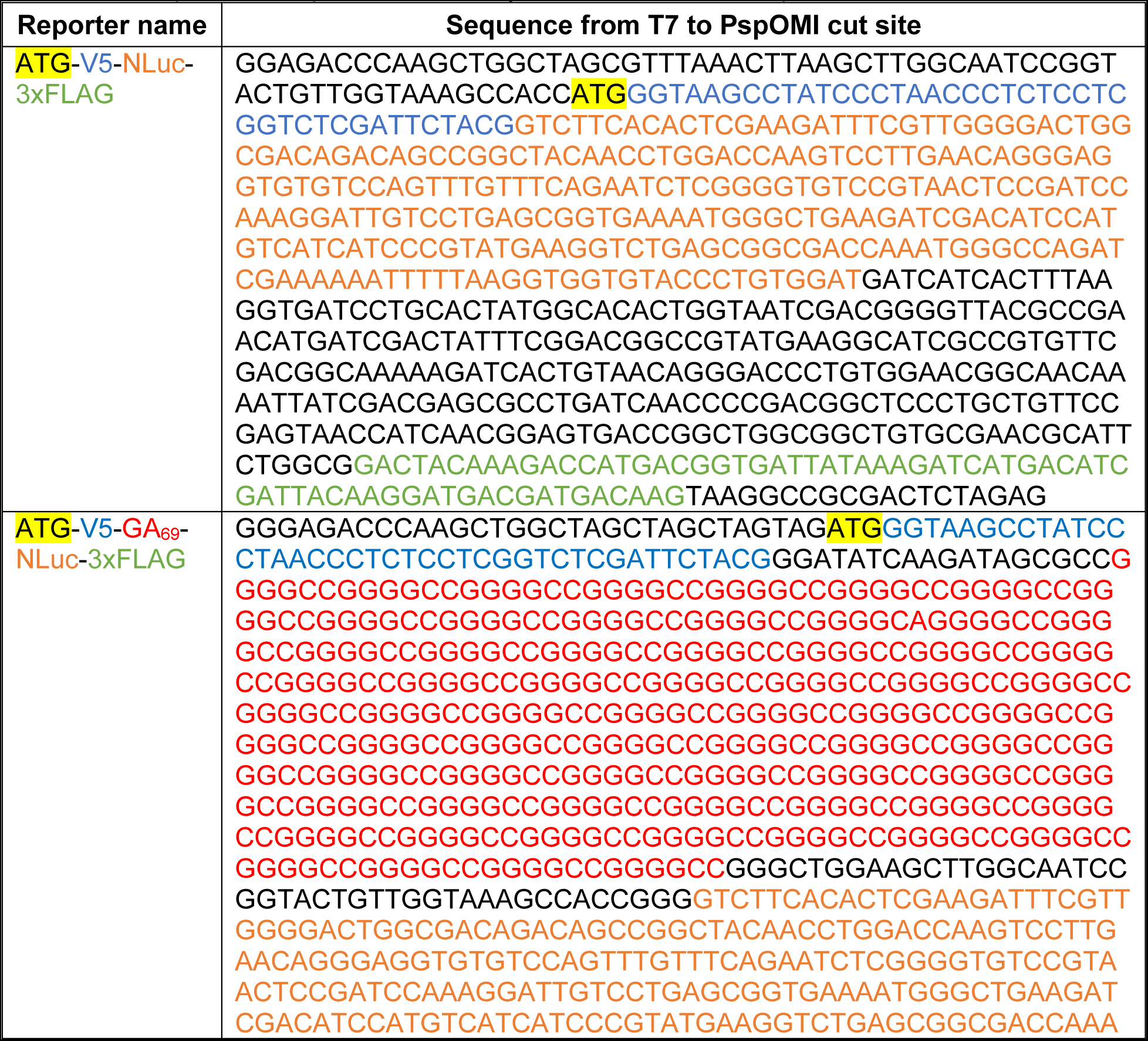
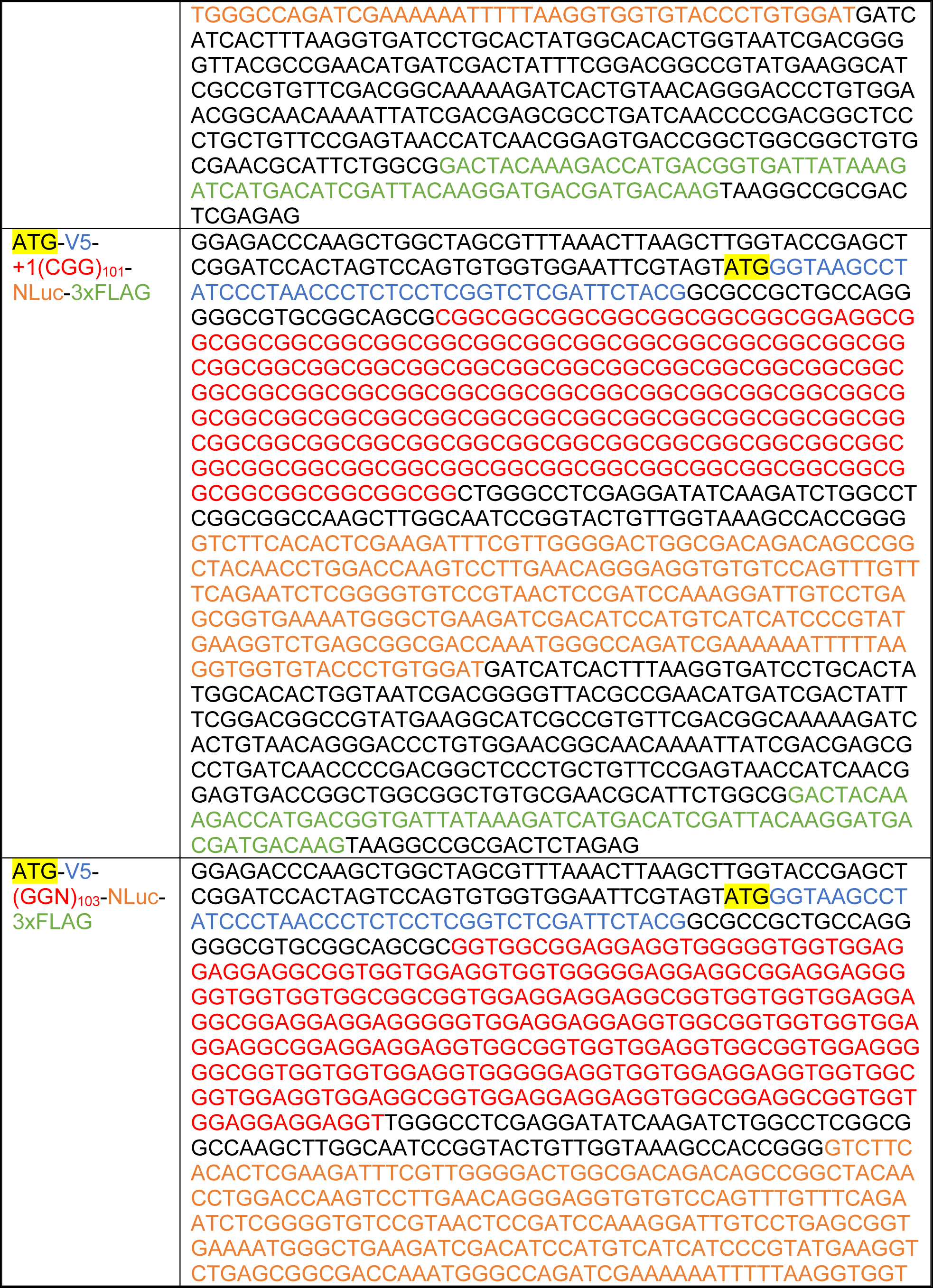
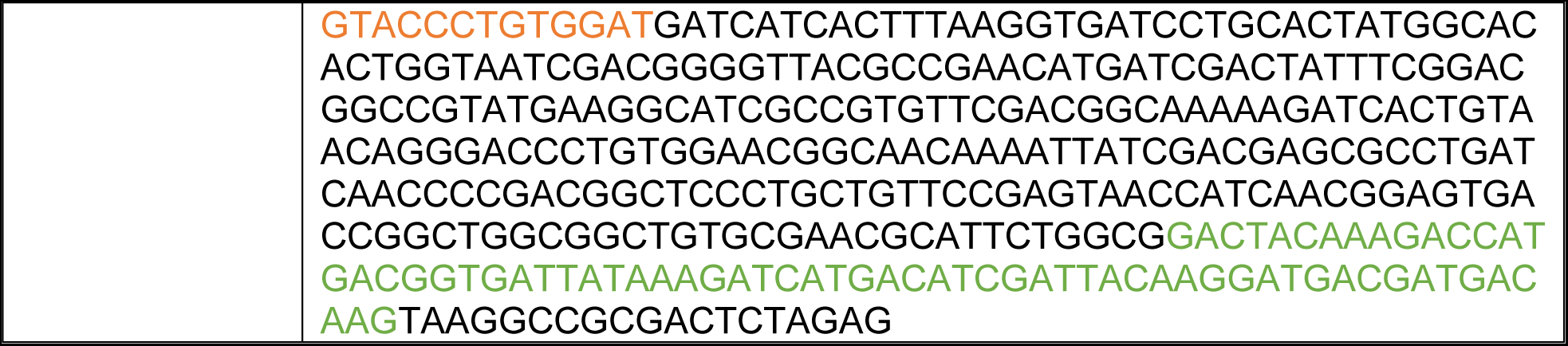
DNA plasmid sequence used to synthesize RNA transcript

**Table 4.**
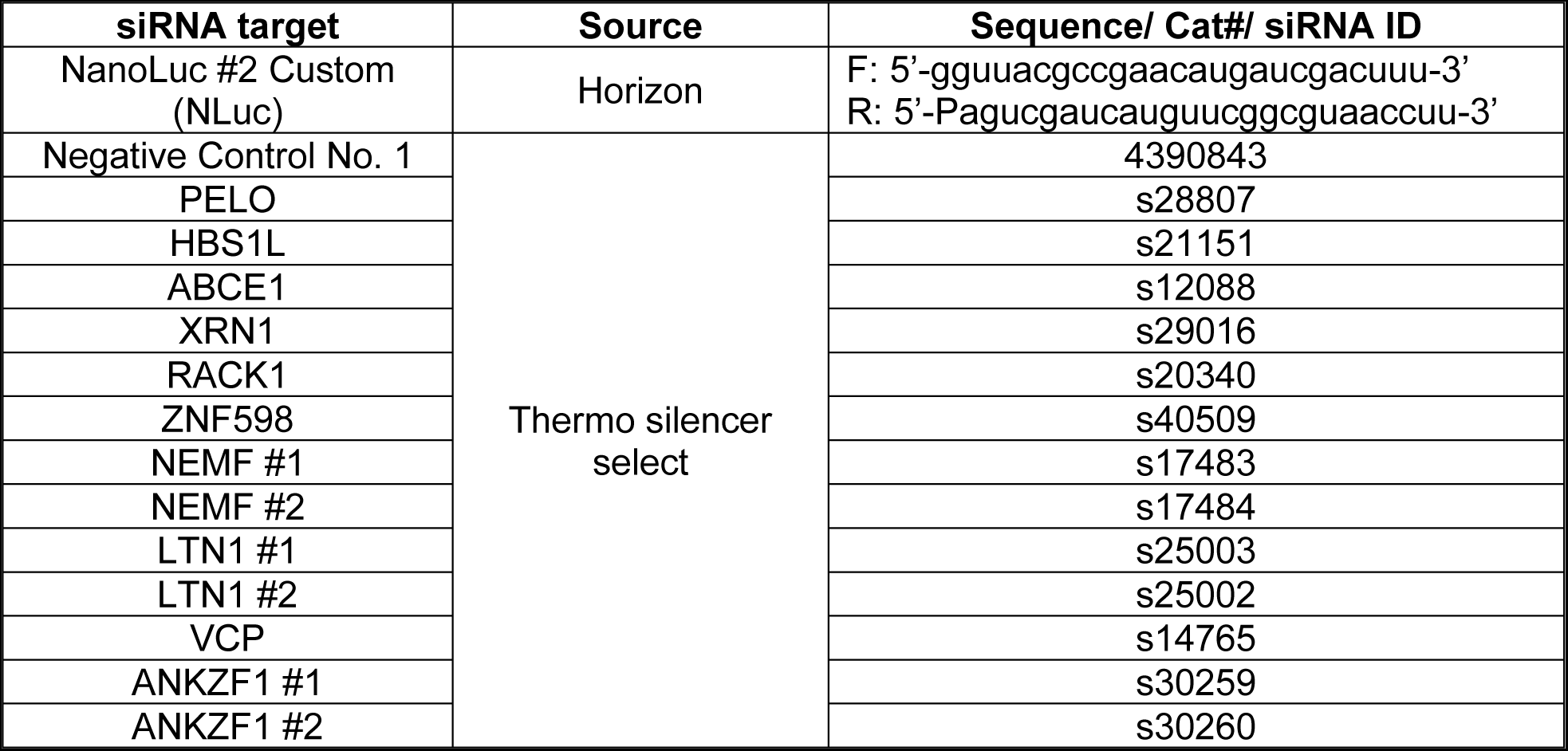
siRNA information

**Table 5.**
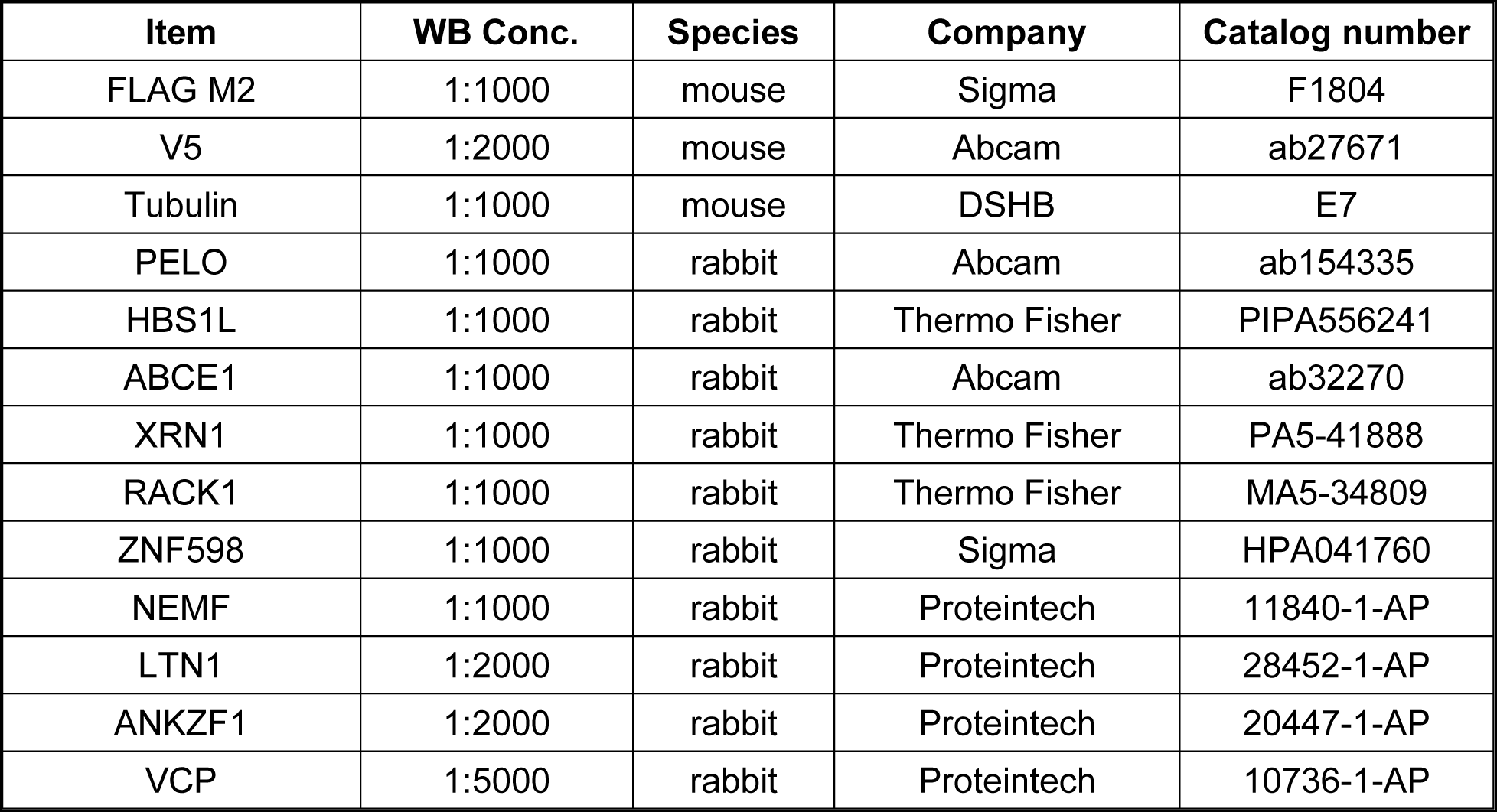
Antibody information

| Item | WB Conc. | Species | Company | Catalog number |
| --- | --- | --- | --- | --- |
| FLAG M2 | 1:1000 | mouse | Sigma | F1804 |
| V5 | 1:2000 | mouse | Abcam | ab27671 |
| Tubulin | 1:1000 | mouse | DSHB | E7 |
| PELO | 1:1000 | rabbit | Abcam | ab154335 |
| HBS1L | 1:1000 | rabbit | Thermo Fisher | PIPA556241 |
| ABCE1 | 1:1000 | rabbit | Abcam | ab32270 |
| XRN1 | 1:1000 | rabbit | Thermo Fisher | PA5-41888 |
| RACK1 | 1:1000 | rabbit | Thermo Fisher | MA5-34809 |
| ZNF598 | 1:1000 | rabbit | Sigma | HPA041760 |
| NEMF | 1:1000 | rabbit | Proteintech | 11840-1-AP |
| LTN1 | 1:2000 | rabbit | Proteintech | 28452-1-AP |
| ANKZF1 | 1:2000 | rabbit | Proteintech | 20447-1-AP |
| VCP | 1:5000 | rabbit | Proteintech | 10736-1-AP |

### Luminescent assays

From a 96-well plate, cells were lysed with 60 μL Glo Lysis Buffer (Promega, E2661) for 5 min at room temperature with constant rocking. NanoGlo Substrate (Promega, N113B) was freshly diluted 1:50 in NanoGlo Buffer (Promega, N112A). For a nano-luciferase assay, 25 μL of the diluted NanoGlo substrate was added to 25 μL of cell lysate. To test cell viability, 25 μL of CellTiter-Glo (Promega, G7573) was added to another 25 μL of lysate. Both assays were mixed for 5 min in a covered opaque 96-well plate then luminescence was measured on a GloMax 96 Microplate Luminometer.

### Immunoblotting

For HEK293, each well of cells from 24-well plates was lysed with 120 μL of RIPA buffer with protease inhibitor (Roche, 11836153001). All protein lysates were denatured with 12% β-mercapto-ethanol in 6x SDS-loading dye, boiled at 95°C for 5 min. 20 μL lysate for each sample was loaded per well on 12% sodium dodecyl sulfate-polyacrylamide gels. To detect stalling products between 10-15 kDa, those samples were loaded on 15% sodium dodecyl sulfate-polyacrylamide gels. Gels were then transferred to PVDF membranes either overnight at 40 V at 4°C, or for 2.5 hr at 320 mAmps in the ice bucket at 4°C. Membranes were blocked with 5% non-fat dry milk for 30-60 min, and all antibodies were diluted in 5% non-fat dry milk. All primary antibody information and probing conditions are listed in **Table 5**. Washes were performed with 1x TBST 3 times, 5 min each. Horseradish peroxidase secondary antibodies were applied at 1:10,000, for 2 hr at room temperature. Bands were then visualized on film. Mild stripping (1.5% Glycine, 0.1% SDS, 1% Tween 20, pH to 2.2 with HCl) was performed with two 10 min incubations at room temperature.

### Immunoprecipitation

Cells were lysed with NP40 lysis buffer (50 mM HEPES-KOH, 150 mM KCl, 0.5% NP40, 0.5 mM DTT, 2 mM EDTA, 1 mM PMSF, protease inhibitors) on ice and incubated at 4°C for 30 min. Lysates were cleared by centrifugation at 20,000 x g for 10 min at 4°C, and the supernatant was transferred into a new tube. Protein concentration for each sample was measured by BCA assay (Thermo Fisher, 23227). 1 mg of total protein was used for each immunoprecipitation. Each lysate was first incubated with 20 μL pre-washed FLAG M2magnetic beads (Sigma, M8823), rotating at 4°C for 2 hr. Next, flow-through was collected and incubated with pre-washed 20 μL of V5-Trap magnetic beads (Proteintech, v5tma) rotating at 4°C for 2 hr. 10% of input and 20% of the supernatant from each step was saved for immunoblot. Afterward, FLAG and V5 beads were washed with NP-40 lysis buffer until the absorbance of the wash supernatant at 280 nm is below 0.05. After the final wash, beads were resuspended with 2x SDS dye and boiled at 95°C for 5 min. The supernatant was collected for immunoblot.

### Drosophila studies

Drosophila were crossed and maintained at 25°C on SY10 food supplemented with dry yeast. For eye phenotyping at a higher temperature, flies were crossed and maintained at 29°C. To measure G4C2 repeat RNA toxicity in flies, a previously characterized GMR-GAL4-driven UAS-(GGGGCC)_28_-EGFP reporter containing fly was used (102). NEMF knockdown flies were obtained from Bloomington Drosophila Stock Center (BDSC) with stock numbers BDSC 36955 and BDSC 25214. Rough eye phenotyping was performed as described earlier (103). In brief, 5-6 virgin female flies expressing the GMR-GAL4-driven UAS-(GGGGCC)_28_-EGFP transgene were crossed with male flies carrying a germline mutation (insertion/disruption) of the fly homolog of NEMF gene (Dmel\Clbn). The rough eye phenotype of flies in F1 progenies was determined at 1-2 days post-eclosion. Rough eye scores were given based on the following eye abnormalities: orientation of bristles, presence of supernumerary bristles, ommatidial fusion, and disarray, presence of necrosis, and shrinkage of the whole eye. Eye images were captured using a Leica M125 stereomicroscope with a Leica DFC425 digital camera. Eye images were scored and analyzed with ImageJ in a blinded manner.

### Maintenance of iPSCs and differentiation into forebrain-like neurons (iNs)

C9 patient-derived induced pluripotent stem cells (iPSCs; CS52iALS-C9n6) and isogenic controls (CS52iALS-C9n6.ISOC3) were obtained from the Cedars-Sinai iPSC Core. iPSCs were maintained in TeSR-E8 media (Stemcell Technologies, 05990) on vitronectin-coated plates, and passaged every 4-5 days using EDTA as described in Weskamp et al. (104). A doxycycline-inducible cassette for induced expression of Ngn1/2 was integrated into the *CLYBL* safe harbor locus of each line, as per Weskamp et al, enabling directed differentiation into forebrain-like glutamatergic iNeurons (iNs) (104–107). Neural progenitor cells (NPCs) were frozen on day 2 of differentiation and stored in liquid N2 until needed. One day before plating NPCs (1x10^6^ cells/well in a 6-well plate), each well was coated overnight at 37°C with 1 mL of 100 μg/mL poly-L-ornithine hydrobromide (PLO, Sigma, P3655) prepared in filter sterilized 0.1 M borate buffer (Fisher Chemical, A73-500) at pH 8.4. PLO was removed by washing with sterile water 3x and air-dried for at least 1 hr. The remainder of the iNs differentiation procedure was as described in Weskamp et al. (104).

### Lentivirus Transduction

Lentiviral shRNA plasmids against NEMF, LTN1, and ANKZF1 were purchased from Horizon Discovery. Lentiviral overexpression of NEMF and ANKZF1 was purchased from VectorBuilder (**Table 6**). Lentiviral overexpression of LTN1 was cloned with NEBuilder Gibson Assembly. In brief, the RPL22 sequence from the plasmid pLV-Ef1a-RPL22-3XHA-P2A-EGFP-T2A-Puro (gift from Hemali Phatnani, Addgene plasmid # 170317) was removed by restriction enzymes digestion with EcoRI-HF and AgeI-HF. The human LTN1 was amplified from pCMV-3TAG8li-hRNF160-3xFLAG with primers (forward: 5’-tttgccgccagaacacaggaccggttaatctgcgctgccaccatgggcggz-3’, Reverse 5’-aattcgtggcgccagatccgggctcgacatcgatgaaaaacg-3’) designed using NEBuilder Assembly Tool v2.7.1. pLV-Ef1a-hLTN1-3XHA-P2A-EGFP-T2A-Puro was generated by Gibson cloning with the hLTN1 fragment generated as described above. Lentiviruses were packed at the University of Michigan Vector Core with HIV lentivirus and then 10x concentrated in 10 mL of DMEM. A GFP vector control was purchased from the University of Michigan Vector Core. Transduction efficiencies as measured by GFP fluorescence were tested in HEK293 cells with 10 μg/mL polybrene following the lentiviral transduction protocol provided by the Vector Core. Knockdown and overexpression of the gene were confirmed with immunoblot. C9 and its isogenic control iNeurons were transduced with lentivirus on Day 6 post differentiation. Cell media were replaced with fresh B27 media on Day 8, and cells were harvested on Day 14.

**Table 6.** Lentiviral constructs

| Plasmid | Source | Cat# |
| --- | --- | --- |
| Lenti-EV-GFP-VSVG | UM Vector Core | UMICHVC-YT-4612 |
| SMARTvector Lentiviral Human NEMF hEF1a-TurboGFP shRNA | Horizon | V3SH11240-224809840 |
| SMARTvector Lentiviral Human LTN1 hEF1a-TurboGFP shRNA | Horizon | V3SH11240-226558312 |
| SMARTvector Lentiviral Human ANKZF1 hEF1a-TurboGFP shRNA | Horizon | V3SH11240-224939332 |
| pLV-EGFP-EF1A-hNEMF [NM_004713.6] | VectorBuilder | VB221216-1307gkc |
| pLV-EF1A-hLTN1-3XHA-P2A-EGFP-T2A-Puro | This paper | N/A |
| pLV-EGFP-EF1A-hANKZF1 [NM_001042410.2] | VectorBuilder | VB900124-1663tuj |

**Table 7.**
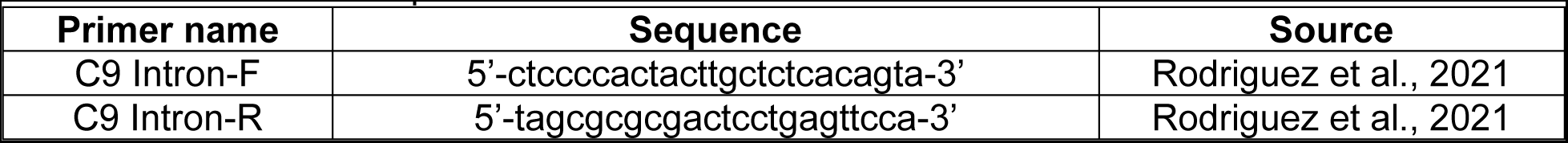
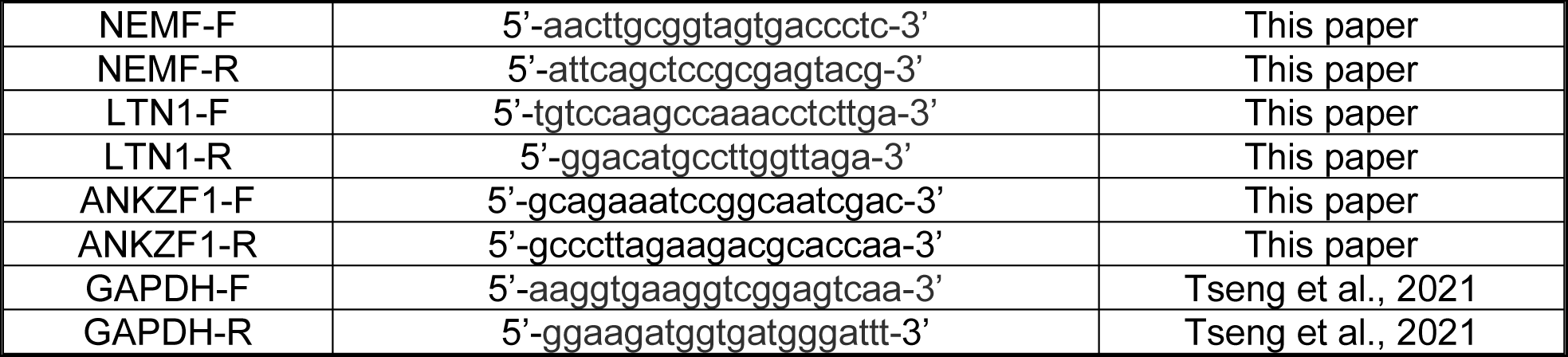
Primer sets for qRT-PCR

## GP MSD

From a 6-well plate, cells were washed with 1x PBS on ice and then harvested by scraping with 200 μL of Co-IP buffer (50 mM Tris-HCl, 300 mM NaCl, 5 mM EDTA, 0.1% triton-X 100, 2% SDS, protease inhibitors, phosphoSTOP) on ice. Lysates were passed through a 28.5G syringe 10 times, spun at 16,000 x g for 20 min at 15°C, and then the supernatant was collected. Levels of polyGP proteins in cell lysates were measured using the Meso Scale Discovery (MSD) electrochemiluminescence detection technology as previously described (108). Briefly, a purified mouse monoclonal polyGP antibody was used as both the capture and detection antibody (Target ALS Foundation, TALS 828.179). For capture, the antibody was biotinylated and used to coat a 96-well MSD Small Spot Streptavidin plate (Meso Scale Discovery, L45SA-2), whereas the detection antibody was tagged with an electro-chemiluminescent label (MSD GOLD SULFO-TAG). An equal amount of each lysate was diluted in TBS and tested in duplicate in a blinded fashion. For each well, the intensity of emitted light, which is reflective of GP abundance and presented as arbitrary units, was acquired upon electrochemical stimulation of the plate.

### Quantitative real-time reverse transcription PCR (qRT-PCR)

RNA from iN lysates was isolated and collected using Quick-RNA MiniPrep Kit (Zymo Research, R1054). 2 μg of RNA per sample was treated with 2 U of TURBO DNase (Thermo Fisher Scientific, AM2238) for 30 min at 37°C twice to remove contaminating genomic and plasmid DNA, and then recovered using the RNA Clean & Concentrator-5 Kit (Zymo Research, R1015). cDNA from each sample was generated from 250 ng of RNA from the previous step with a mixture of oligo(dT) and random hexamer primers (iScript cDNA Synthesis Kit, Bio-Rad, 1708891). Finally, cDNA abundance was measured using iQ SYBR Green Supermix (Bio-Rad, 1708882) from an iQ5 qPCR system (Bio-Rad), and the appropriate primers at 100 nM. Primer information is listed in **Table 7**. cDNA abundance was quantified using a modified ΔΔCt method recommended by the manufacturer.

### Polysome profiling

HEK293 cells at 85-95% confluency in a 15 cm culture dish were treated with 100 μg/mL cycloheximide (CHX) for 5 min at 37°C. The culture dish was placed on ice, washed with 5 mL ice-cold PBS containing 100 μg/mL CHX, harvested by scraping with another 5 mL cold PBS + CHX, and centrifuged at 1200 x g at 4°C for 5 min. PBS was aspirated and the pellet was resuspended in polysome-profiling lysis buffer (20 mM Tris-HCl (pH 7.5), 150 mM NaCl, 15 mM MgCl2, 8% (vol/vol) glycerol, 20 U/mL SUPERase, 80 U/ml murine RNase inhibitor, 0.1 mg/mL heparin, 100 μg/mL CHX, 1 mM DTT,1x EDTA-free protease inhibitor cocktail, 20 U/mL Turbo DNase, 1% Triton X-100). Lysates were vortexed for 30 sec, passed through a 21G needle 10 times, and incubated on ice for 5 min. Cellular debris was pelleted at 14,000 x g at 4°C for 10 min, and the supernatant was transferred to a pre-cooled tube. Total lysate RNA was estimated by NanoDrop. Lysates were flash-frozen in liquid nitrogen and stored at -80°C until fractionation.

Sucrose gradients were prepared by sequentially freezing 2.7 mL of 50, 36.7, 23.3, and 10% sucrose (wt/vol) in 13.2 mL thin wall polypropylene tubes (Beckman Coulter, 331372). Sucrose-gradient buffer consisted of 20 mM Tris-HCl (pH 7.5), 150 mM NaCl, 15 mM MgCl2, 10 U/mL SUPERase, 20 U/mL murine RNase inhibitor, 100 μg/mL CHX, and 1 mM DTT. Before use, gradients were thawed and linearized overnight at 4°C. For fractionation, approximately 50 μg total RNA was applied to the top of the sucrose gradient. Gradients were spun at 35,000 rpm at 4°C for 3 hr using a Beckman Coulter Optima L-90 K ultracentrifuge and SW 41 Ti swinging-bucket rotor. Gradients were fractionated with Brandel’s Gradient Fractionation System, measuring absorbance at 254 nm. The detector was baselined with 60% sucrose chase solution and its sensitivity was set to 0.05. For fractionation, 60% sucrose was pumped at a rate of 1.5 mL/min. Brandel’s PeakChart software was used to collect the profile data.

### Data analysis

Prediction of RNA structure and calculation of the minimum free energy was computed by The Vienna RNA Websuite, RNAfold 2.5.1. (109). Statistical analyses were performed with GraphPad Prism9.5.1. All luciferase activity was calculated by normalizing the nano-luciferase signal with cell titer. For comparison of NLuc reporter luciferase activity assays, GP MSD response, and fly eye phenotype quantification, we used two-tailed unpaired Student’s t-tests with Welch’s correction for multiple comparisons to confirm the statistical difference between control and multiple experimental groups within each sample. To assess group effects, we used a two-way ANOVA with Sidak’s multiple comparison tests to compare the differences between groups within different samples. Fly eye necrosis and width measurements were done with reviewer genotype-blinded analysis by at least two independent investigators to avoid subjective bias. Fly eye width measurement was performed with ImageJ (www.imagej.nih.gov/ij/). Experiments were performed with a minimum of three independent biological samples (n > 3) with technical replication of results from each sample. Fly experiments were done using multiple crosses with a minimum of 10 flies analyzed per group per cross. Further statistical analysis details are included in figure legends including the numbers of analyzed samples, statistical tests, and P values.

## RESULTS

### RQC pathway factor depletion enhances RAN translation from GC-rich repeat sequences

Slowing or stalling of translational elongation can result in ribosomal collisions and engagement of ribosomal quality control pathways. He hypothesized that GC-rich repetitive elements would be prone to such events. We therefore performed a targeted modifier screen at two disease-associated repetitive elements (G4C2 hexanucleotide or CGG trinucleotide repeats) for factors involved in mRNA and protein surveillance pathways to evaluate their role in regulating RAN translation **(Figure 1A)**. Specifically, we assessed the impact of lowering expression of RNA degradation pathway factors PELO, HBS1L, ABCE1, and XRN1; ribosome stalling sensing factors RACK1 and ZNF598, or RQC pathway factors NEMF, LTN1, VCP, and ANKZF1. Our group previously generated a series of RAN translation-specific nano-luciferase reporters for both G4C2 repeats and CGG repeats **(Figure 1B)** (44, 49). We used those well-characterized reporters for the screen. Validation of gene knockdown for each factor was confirmed by immunoblotting **(Figure S1)**. For factors involved in Ribosome associated RNA degradation, depletion of XRN1 increased RAN translation at both G4C2 and CGG repeats, while depletion of HSB1L selectively increased RAN translation from CGG repeats **(Figure 1C)**. However, the largest effect was observed for factors involved in the RQC pathway that classically engages the ribosomes after stall detection and ribosome splitting. Depletion of NEMF, LTN1, or ANKZF1 markedly and selectively increased the accumulation of RAN translation products from both G4C2 and CGG repeats, with increases between 2 and 5-fold compared to a non-targeting siRNA control. We, therefore, focused our attention on these RQC pathway factors and their effect on RAN translation.

**Figure 1.**
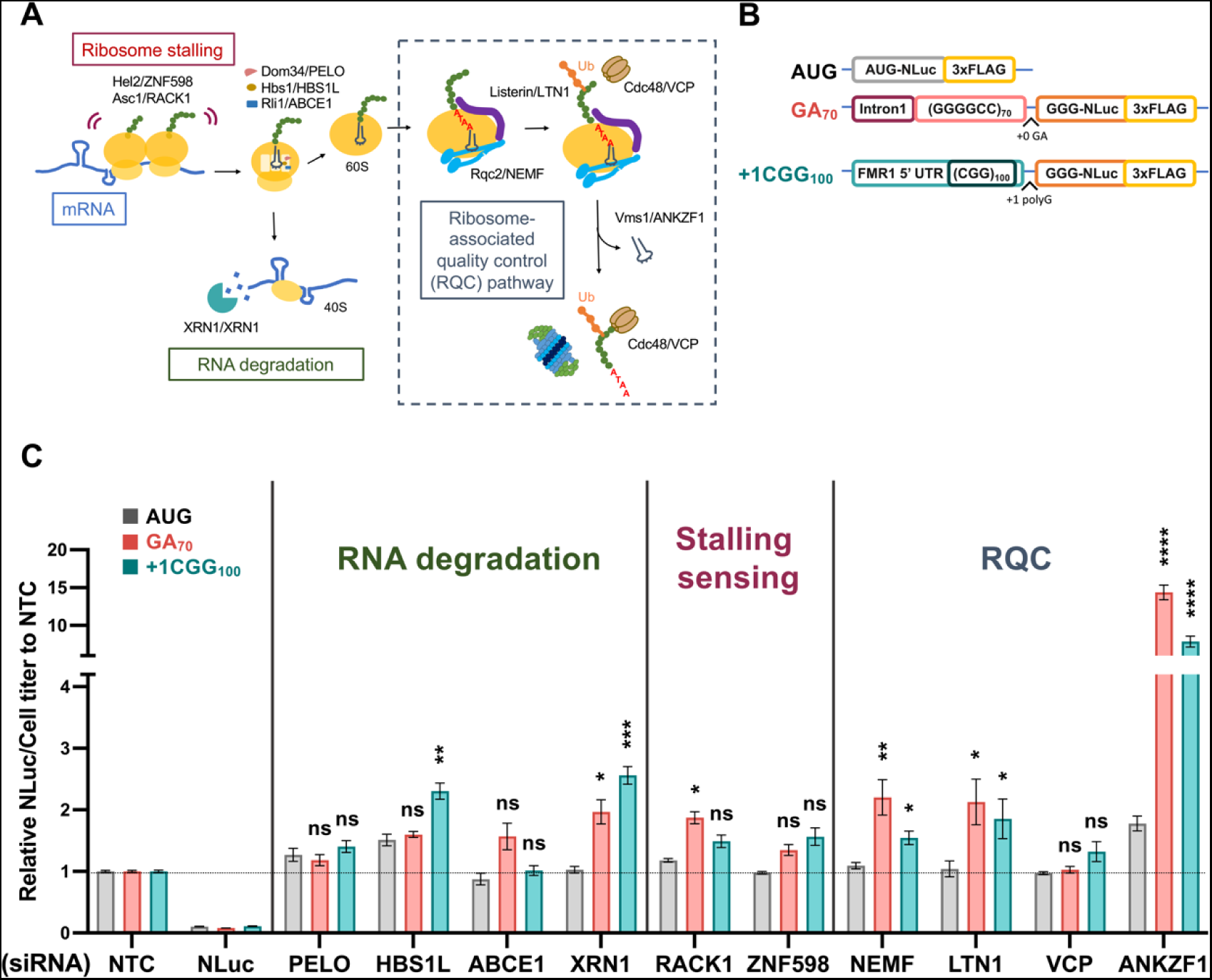
NEMF, LTN1, and ANKZF1 act as genetic modifiers of RAN translation from G4C2 and CGG repeats. (**A**) Schematic of mRNA and protein surveillance pathways(63).(**B**) Schematic of luciferase reporters used to assess RAN translation product abundance. (**C**) Targeted screen of factors involved in mRNA and protein surveillance pathways. The relative expression of NLuc to cell titer was compared to a non-targeting siRNA control (NTC). Data represent mean with error bars ± SEM of n = 6 cross at least 2 independent experiments ns = not significant; \**P* < 0.05; \*\**P* ≤ 0.01; \*\*\**P* ≤ 0.001; \*\*\*\**P* ≤ 0.0001, multiple unpaired t-test after Welch’s correction.

### NEMF, LTN1, and ANKZF1 effects on RAN translation are repeat length-dependent

Previous studies found that knockdown of NEMF and LTN1 can increase the production of GR and PR DPRs from non-GC-rich sequences (98, 100). This difference was thought to result from the positively charged arginine residues in PR and GR DPR tracks, which can interact with the ribosome exit tunnel to elicit ribosomal stalling (97, 99, 110). However, RNA secondary structures can also elicit ribosomal stalling (111–113). As GC-rich repeats form stable secondary structures and the RNA helicases that resolve these structures are implicated in RAN translation (114–117), we hypothesized that ribosome stalling would occur during RAN translation from GC-rich repeats, leading to RQC pathway activation. To assess this, we first validated the effects of NEMF, LTN1, and ANKZF1 on RAN translation with a second set of siRNAs. Knockdown of NEMF, LTN1, or ANKZF1 with this second set of siRNAs elicited similar increases in GA DPR RAN production from G4C2 and CGG repeats **(Figure 2, Figure S2A-C)**. We next assessed whether this impact by RQC factors was dependent on the length of the G4C2 repeat. At 35 G4C2 repeats, we did not see any effect of NEMF knockdown on GA frame RAN translation, suggesting repeat-length dependence **(Figure 2A, 2B)**. Similarly, the depletion of LTN1 significantly increased RAN translation from both GA35 and GA70 repeat reporters, but the effect on the larger repeat was greater **(Figure 2C)**. As NEMF helps recruit LTN1 to assemble the RQC complex, we asked if there is a synergetic effect of NEMF and LTN1 knockdown on RAN translation. When we performed the knockdown of both NEMF and LTN1 together, we observed a mild additive effect, with slightly more RAN translation production from GA35, GA70, GP70, and GR70 reporters **(Figure 2D)**. Similarly, the depletion of ANKZF1 had a modest effect on translation from an AUG reporter with no repeat and from our GA35 reporter, but a larger effect on product generation from the GA70 reporter **(Figure 2E)**.

**Figure 2.**
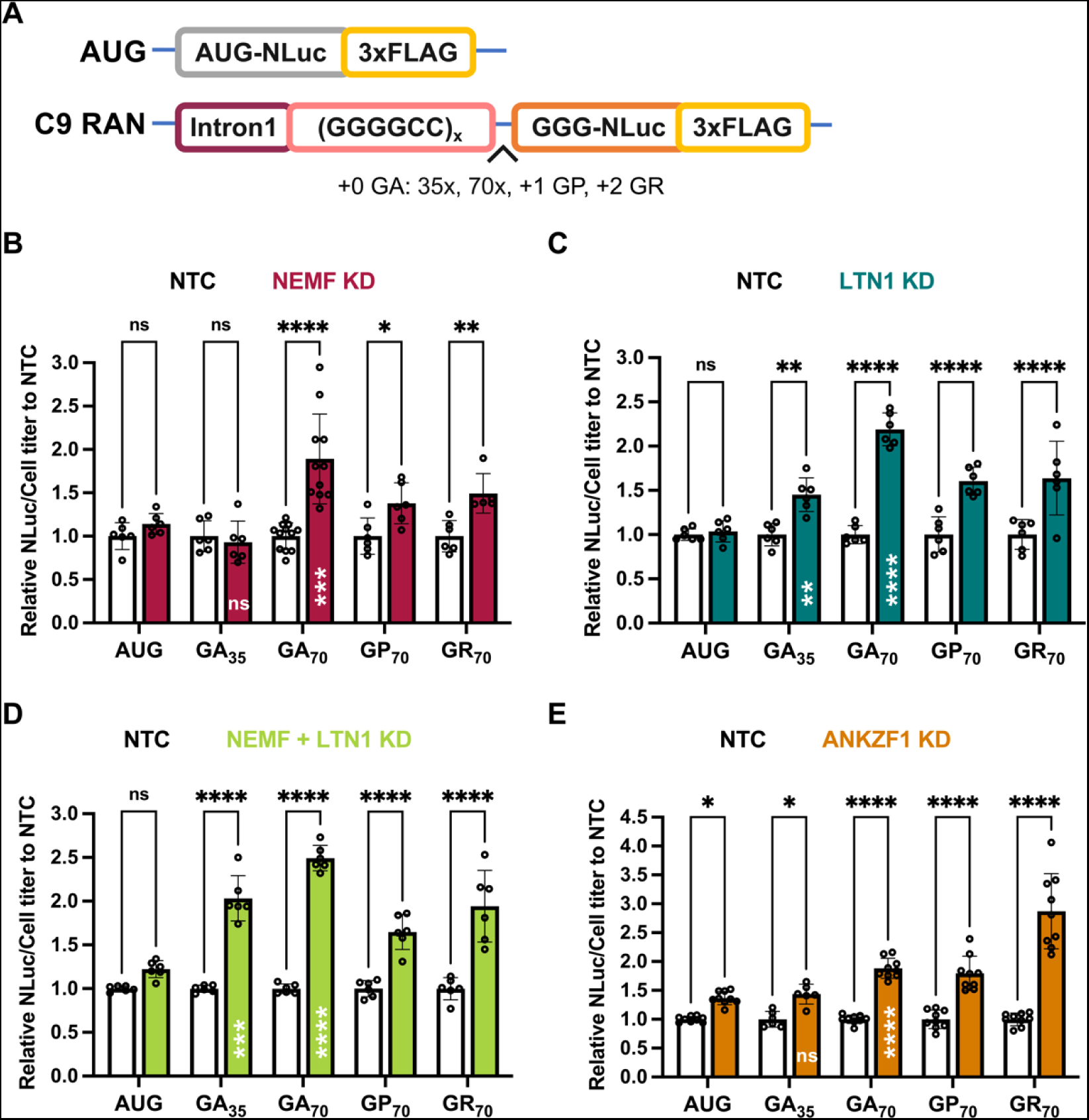
Depletion of NEMF, LTN1, and ANKZF1 enhances G4C2 C9 RAN translation in a repeat length-dependent manner across all reading frames. (**A**) Schematic AUG-driven NLuc-3xFLAG and C9 RAN G4C2 repeat length and reading frame reporters. (**B-E**) Luciferase assays after NEMF, LTN1, both NEMF and LTN1 or ANKZF1 depletion. All graphs show mean with error bars ± SD. Each N is shown as an open circle (n=6-9/group). Asterisks above each bar are comparisons of expression between NTC and gene(s) knockdown. ns = not significant; \**P* < 0.05; \*\**P* ≤ 0.01; \*\*\**P* ≤ 0.001; \*\*\*\**P* ≤ 0.0001, as determined with two-way ANOVA with Sidak’s multiple comparison test. Asterisks placed inside each bar are comparisons between AUG-driven no-repeat control and different repeat lengths of the GA frame of cells treated with gene(s) knockdown. ns = not significant; \*\**P* ≤ 0.01; \*\*\**P* ≤ 0.001; \*\*\*\**P* ≤ 0.0001, represent unpaired t-tests with Welch’s correction for multiple comparisons.

If RNA secondary structure contributes to ribosomal stalling, then the effect of knockdown of RQC factors should influence RAN translation production from all 3 potential reading frames. To address this, we utilized (G4C2)70 RAN translation reporters, where the NLuc tag was in the GP (+1) or GR (+2) reading frames **(Figure 2A)**. Knockdown of NEMF increased RAN from (G4C2)70 sequences in all 3 reading frames **(Figure 2B)**. Similarly, the knockdown of LTN1 increased RAN product accumulation from (G4C2)70 sequences in GA, GP, and GR frames **(Figure 2C)**. When we knocked down both NEMF and LTN1 together, we observed a mild additive effect, with more production from GA35, GA70, GP70, and GR70 RAN reporters compared to controls **(Figure 2D)**. This suggests that NEMF and LTN1 are functioning on the same pathway in RAN translation. Knockdown of ANKZF1 significantly increased RAN production from (G4C2)70 sequences in all three reading frames, with greater effects on the GR frame **(Figure 2E)**. Taken together, these data suggest that depleting RQC factors increase RAN translation in a repeat length-dependent but repeat reading-frame independent fashion.

### Detection of short/truncated translation products from G4C2 and CGG transcripts

RQC pathway activation typically triggers the degradation of partially made translation products through a proteosome-dependent process. Prior studies utilized a dual fluorescent tagging system with GFP and RFP bracketing a ribosome-stall inducing sequence coupled with flow cytometry to measure stalling during translation (91). In this set up ‘stalling’ events are measured by detecting the signal of one (stalled/truncated) or both reporters (full-length). However, such truncated products would not be detectable by either carboxyl-terminal nano-luciferase (NLuc) signal or FLAG tag western blots. We, therefore, generated a new set of reporter constructs with an AUG-initiated V5 tag at the amino-terminus of 69 G4C2 repeats in the GA frame or 100 CGG in the polyG frame. For each construct, we retained an in-frame NLuc and 3xFLAG at the carboxyl-terminus **(Figure 3A)**. To avoid the generation of potentially aberrant RNA products from plasmids that might complicate our data interpretation, we transfected in vitro synthesized mRNAs with either G4C2 repeats or CGG repeats into HEK293 cells. We then performed a dual IP experiment where we first used FLAG magnetic beads to pull down the full-length products and then performed a second IP with V5 on the flowthrough to enrich for partially made products that only have the amino terminus of the protein **(Figure 3B)**. We detected partially made products, approximately 10-15 kDa in size, only with the V5 antibody for both GA **(Figure 3C)** and polyG **(Figure 3D)**. In contrast, constructs lacking the GC-rich repeats did not generate any partially made products **(Figure S3A-C)**. These data suggest that ribosome stalling presumably occurs during translational elongation from GC-rich repeats - even when the protein products from those repeats lack arginine or similarly charged amino acids.

**Figure 3.**
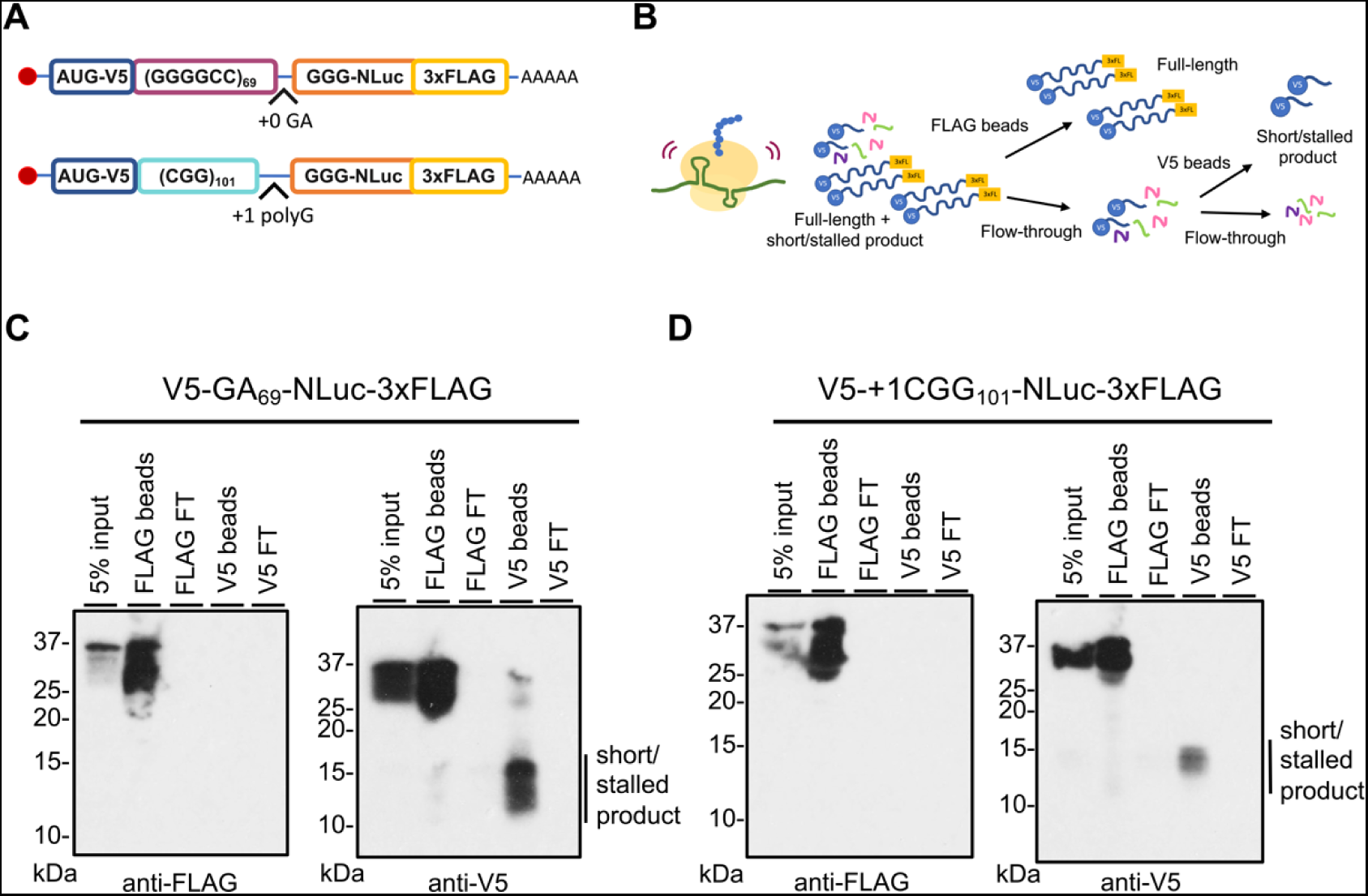
Detection of partially made products translated from GC-rich transcripts. (**A**) Schematic of AUG-V5-(G4C2)_69_-NLuc-3xFLAG in GA frame and AUG-V5-+1(CGG)_101_-NLuc-3xFLAG in polyG frame. (**B**) FLAG and V5 IP workflow to enrich for incomplete products generated from GC-rich transcripts. 3xFL: 3xFLAG (**C-D**) V5 antibody IP reveals short/stalled products (line next to the blot) that are generated from G4C2 (panel C) or CGG repeats (panel D) that do not contain the carboxyl-terminal FLAG tag. FT: Flow-through.

To assess whether NEMF, LTN1, or ANKZF1 might affect the generation or accumulation of these truncated repeat products, we repeated the studies as above after depletion of NEMF + LTN1 or ANKZF1. From immunoblot analysis, we detected an increase in both the abundance of the full-length products and the truncated products after double knockdown of NEMF + LTN1 or after knockdown of ANKZF1 from both (G4C2)69 and (CGG)100 repeats **(Figure 4A-D)**. However, we do not see any change in the expression or generation of products from a no-repeat control **(Figure S3B-C)**. These observations suggest that short/stalled products are generated normally during translation through GC-rich repeats and that their abundance is impacted by RQC factor abundance.

**Figure 4.**
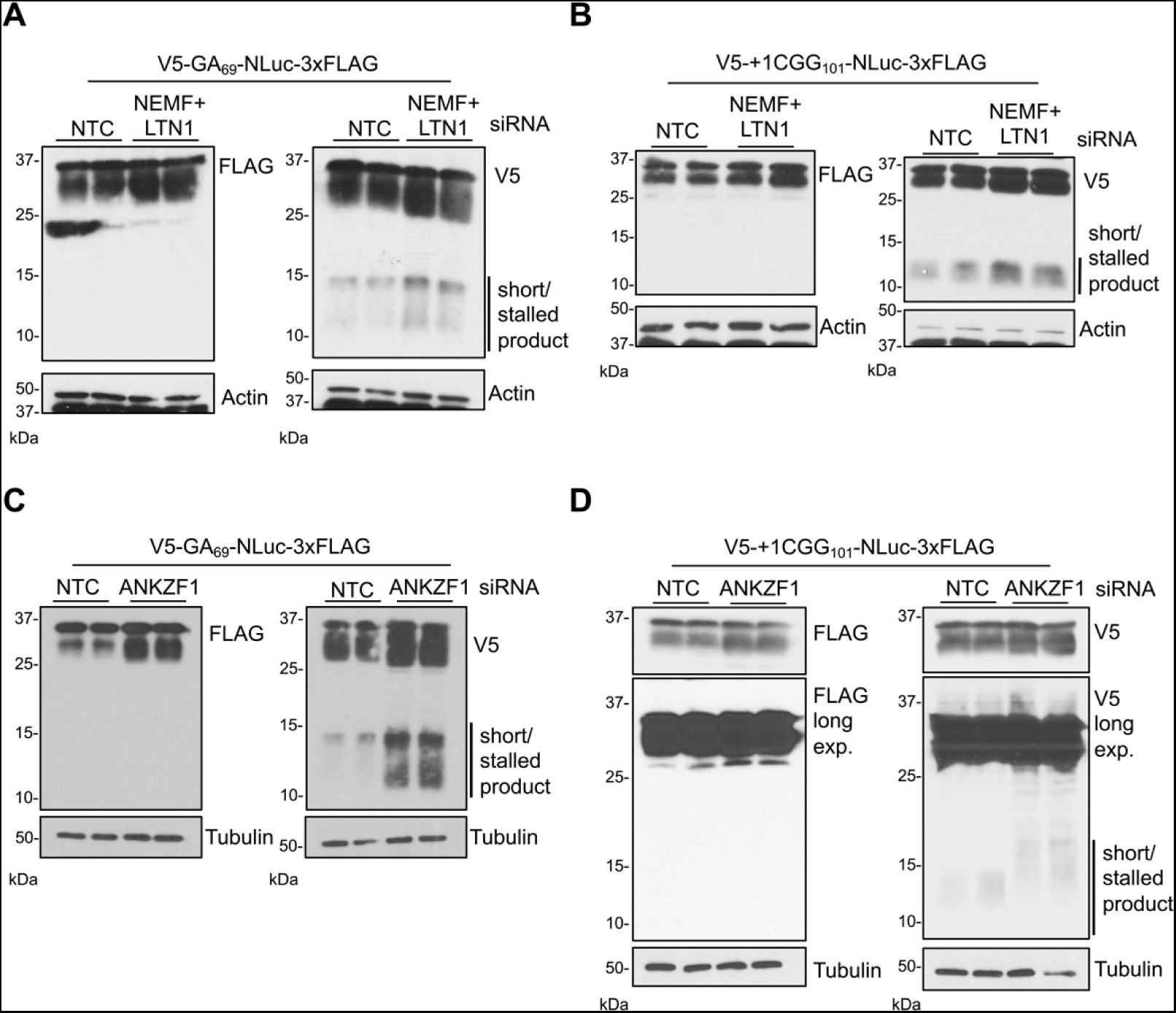
Depletion of NEMF, LTN1, and ANKZF1 enhances the generation of both full-length and partially made GA and polyG products from GC-rich transcripts. (**A-D**) Immunoblots of HEK293 cells transfected with in vitro synthesized G4C2 (GA frame) and CGG repeat RNA reporters (polyG frame) in the presence and absence of NEMF + LTN1, and ANKZF1. Samples were processed as in Figure 3B. Blots represent two biological replicates. Lines next to the blots indicate short/stalled product products.

### Repeat RNA sequence determines the impact of RQC factor depletion

Prior studies are conflicted as to whether GA DPRs generated from non-repetitive mRNA sequences induce ribosomal stalls (98, 100). Our data suggests that the repeat RNA sequence and structure may be sufficient to elicit ribosomal stalling and RQC activation **(Figure 3, 4)**. To assess this more formally, we took advantage of the redundant nature of RNA codons to generate a reporter that would make a polyG product but lack the CGG repeat RNA structure **(Figure 5A-B)**. Glycine is encoded by GGN, where N is any nucleotide. Therefore, we generated constructs with identical AUG initiation codons above either a GGC repeat or a GGN repeat that should not form a strong hairpin structure **(Figures 5A-C)**. Both constructs will generate the same polyG-containing protein that we can measure using NLuc **(Figure 5C)**. Compared to a non-targeting siRNA control, depletion of NEMF, LTN1, or ANKZF1 significantly enhanced the production of the polyG protein from the AUG-V5-(CGG)101 reporter, which suggests that the modulation of these factors is acting primarily at the stage of translational elongation rather than RAN translational initiation. In contrast, depletion of either NEMF or LTN1 had no impact on polyG protein generation from the AUG-V5-(GGN)103 construct or an AUG-V5-NLuc no-repeat control **(Figure 5D)**. Depletion of ANKZF1 did modestly enhance polyG production from the AUG-V5-(GGN)103 construct, but the effect was significantly weaker than that seen for polyG protein production from the AUG-V5-(CGG)101 reporter **(Figure 5D)**. Together, these results suggest that the repeat sequence plays a key role in eliciting ribosomal stalling and the interplay of RQC factors with CGG repeat translation.

**Figure 5.**
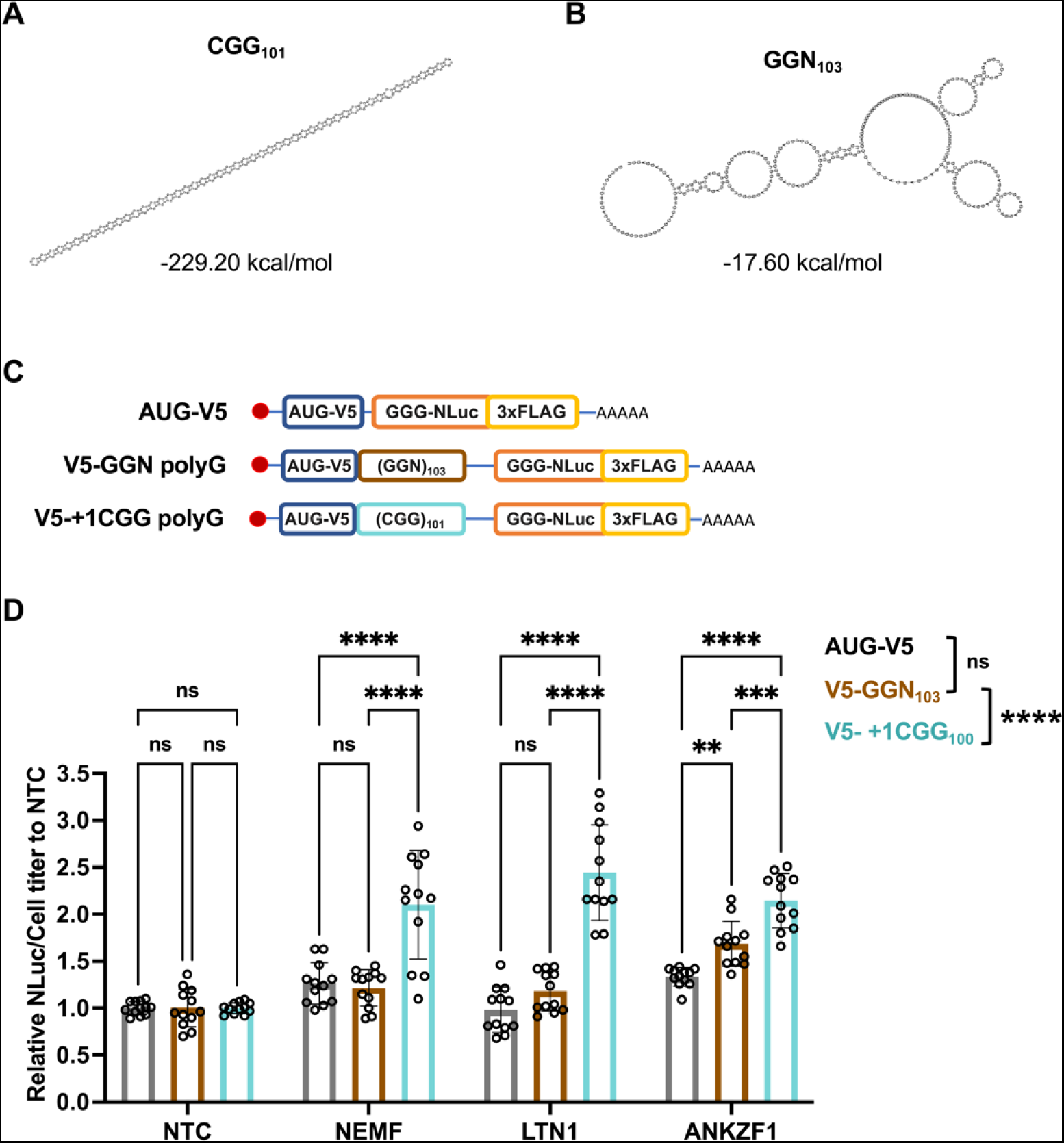
Enhancement of polyG production with NEMF, LTN1, and ANKZF1 depletion requires the CGG repeat RNA structure. (**A-B**) Prediction of the optimal RNA secondary structure and calculation of the minimum free energy in CGG_101_ and GGN_103_ repeats. Only the repeat region from each repeat was used to predict the RNA secondary structure and calculate the minimum free energy. The results were computed by RNAfold 2.5.1. (**C**)Schematics of AUG-V5-NLuc-3xFLAG, AUG-V5-+1(CGG)_101_-NLuc-3xFLAG, and AUG-V5-(GGN)_103_-NLuc-3xFLAG transcripts. (**D**) Knockdown of NEMF, LTN1, and ANKZF1 in HEK293 with no-repeat control, polyG from GGN repeats, and CGG repeats RNA transfection. Data represent means with error bars ±SD of n = 12, ns = not significant; \*\**P* ≤ 0.01; \*\*\**P* ≤ 0.001; \*\*\*\**P* ≤ 0.0001 by Tukey’s multiple comparisons tests. The statistic result placed in the legend is group comparisons by two-way ANOVA Sidak’s multiple comparisons tests.

### Depletion of NEMF and ANKZF1 aggravates toxicity in *Drosophila* model of C9 FTD/ALS

A prior study demonstrated a worsening of rough eye phenotypes with depletion of NEMF in *Drosophila* expressing dipeptide GR or PR from constructs lacking a G4C2 repeat, which again suggested a direct role for serial arginine translation in RQC pathway activation (98, 101). To assess the impact of modulating RQC factor expression on G4C2 repeat-associated phenotypes in vivo, we utilized an established *Drosophila* model of C9ALS/FTD that supports RAN translation from G4C2 repeats and elicits significant toxicity resulting in a rough eye phenotype when expressed using a GMR-GAL4 driver (102). Genetic ablation of NEMF led to a more severe rough eye phenotype compared to the control cross in both NEMF depletion lines **(Figure 6A)**. Rearing the G4C2 repeat-expressing flies at a higher temperature (29°C) led to more severe eye degeneration as marked by physical constriction of eye width (103, 118). Depletion of NEMF mitigated this decrease in eye width **(Figure 6B)**. Together these data suggest that depletion of NEMF enhances G4C2 repeat elicited toxicity in *Drosophila*.

**Figure 6.**
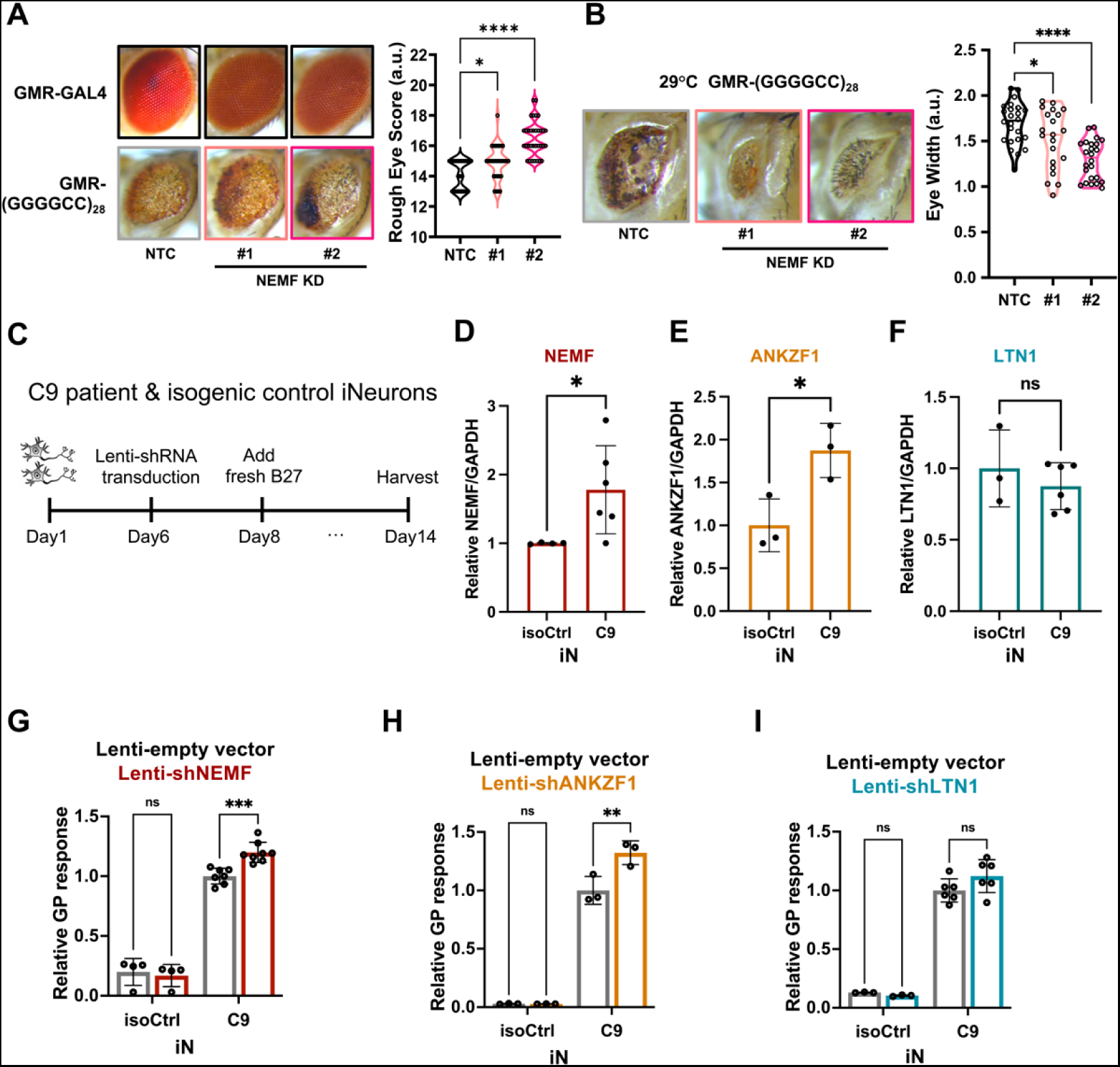
Depletion of NEMF enhances repeat-associated toxicity in a fly model of C9 ALS/FTD and DPR accumulation in human neurons. (**A)** Representative images of *Drosophila* eyes expressing (G4C2)28 repeats under the GMR-GAL4 driver in the presence or absence of NEMF at 25°C [BDSC36955 #1 and BDSC25214 #2]. Rough eye phenotypes quantified using a nominal scoring system are shown as violin plots on the right. Individual flies are represented by single data points. n= 30-33/genotype. (**B**) (G4C2)28 repeats expressed with a GMR-GAL4 driver at 29°C show decreased eye width that is enhanced by the depletion of NEMF, as quantified on the right. Graphs represent the mean with error bars ±SD, n = 21-24. For A and B, \**P* < 0.05; \*\*\*\**P* ≤ 0.0001 by one-way ANOVA with Dunnett’s multiple comparison test. (**C**) Schematic workflow for studies with C9ALS patient-derived iNeurons (iN). (**D-F**) RNA Expression of NEMF, LTN1, and ANKZF1 transcripts from C9ALS and isogenic control iN lysates. Data represent means with error bars ±SD. n = 3-6/gene, ns = not significant; \**P* < 0.05 by Student’s t-test. (**G-I**) Quantification of GP by MSD assay from C9ALS and isogenic control iNs treated with lenti-empty vector or lenti-shRNA of NEMF, LTN1, or ANKZF1. Data represent mean ±SD; n = 3-6, ns = not significant; \*\**P* ≤ 0.01; \*\*\**P* ≤ 0.001 by two-way ANOVA with Sidak’s multiple comparison test.

### Depletion of NEMF and ANKZF1 enhances RAN translation in C9ORF72 human neurons

We next assessed whether modulating RQC factors also influence RAN translation product accumulation from the endogenous repeat locus in *C9ORF72* ALS patient-derived neurons. To this end, we utilized a well-characterized pair of C9 patient iPSC-derived iNeurons (iN) with an accompanied isogenic control line (119). These iN lines contain a doxycycline-inducible Ngn1/2 cassette that allows for rapid neuronal differentiation (106). After two weeks of differentiation, we harvested cell lysates and performed an MSD assay to measure GP DPR product abundance **(Figure 6C)**. In parallel, we performed a qRT-PCR analysis of RNA abundance for different RQC factors. We observed a significant increase of NEMF and ANKZF1 transcript expression in C9ALS iNeurons compared to isogenic controls differentiated in parallel, suggesting that there could be a compensatory upregulation of RQC machinery elicited by the repeat expansion **(Figure 6D, E)**. There was no change in LTN1 transcript abundance between these C9ALS iNeurons and their isogenic control **(Figure 6F)**.

As expected, we were able to reliably measure GP DPR abundance in C9ALS iNeurons but not in isogenic control neurons across multiple differentiations **(Figure 6G-I).** To assess whether altering RQC factor abundance could impact endogenous RAN product abundance, we utilized lentiviral delivery of shRNAs against each factor. In line with our reporter assays, we observed a significant increase in GP DPR abundance following the depletion of NEMF or ANKZF1 in C9 iN compared to iNeurons treated with a lentiviral control **(Figure 6G, H)**. However, the depletion of LTN1 did not significantly impact GP DPR abundance **(Figure 6I)**. The RQC pathway typically pairs with the mRNA surveillance pathway after the separation of 80S ribosomes into 40S and 60S subunits to degrade the template mRNA. Therefore, we assessed whether endogenous C9 RNA abundance might be impacted by the depletion of NEMF, LTN1, and ANKZF1. Using primers that specifically target the first intron of C9RNA that contains the repeat, we observed a modest increase of C9 transcript abundance with depletion of NEMF or LTN1 **(Figure S4A-B)**. However, we observed no effect on the depletion of ANKZF1 compared to a lentiviral control **(Figure S4C)**. Taking our data together with the previous study, whether the activation of RQC degraded the C9 RNA is still unclear (98).

### Enhancing the expression of RQC factors suppresses RAN product accumulation

As depletion of NEMF, LTN1, and ANKZF1 increase GA and polyG product accumulation from GC-rich repeats, we wondered if boosting NEMF, LTN1, or ANKZF1 expression in cells might suppress RAN translation or lower RAN product accumulation. To assess this, we overexpressed each of the RQC factors (NEMF, LTN1, and ANKZF1) in conjunction with the AUG-driven no repeat control, G4C2 GA reporter, or the CGG polyG reporter. Overexpression of NEMF and LTN1 decreased both GA and polyG RAN product accumulations **(Figure 7A-B)**. Overexpression of ANKZF1 decreased the accumulation of RAN products from all three (GA, GP, and GR) reading frames from G4C2 repeats without impacting an AUG-driven no-repeat control **(Figure 7C)**. In contrast, overexpression of ANKZF1 did not significantly decrease polyG RAN product abundance **(Figure 7D)**. Next, we transduced C9 patient iPSC-derived iNeurons (iN) with lentivirus expressing hNEMF, hLTN1, or hANKZF1 and subsequently measured the GP DPR abundance **(Figure 6C)**. Overexpression of either NEMF or ANKZF1 in C9 iN significantly decreased GP abundance compared to a lentiviral control **(Figure 7E, F)**. However, there was no significant increase in GP abundance seen with hLTN1 expression **(Figure 7G)**. Notably, overexpression of NEMF, LTN1, or ANKZF1 did not affect C9 intronic transcript abundance **(Figure S4D-F)**. Taken together, these data suggest modulation of RQC factors can directly impact RAN product accumulation from both reporters and endogenous loci in patient-derived neurons.

**Figure 7.**
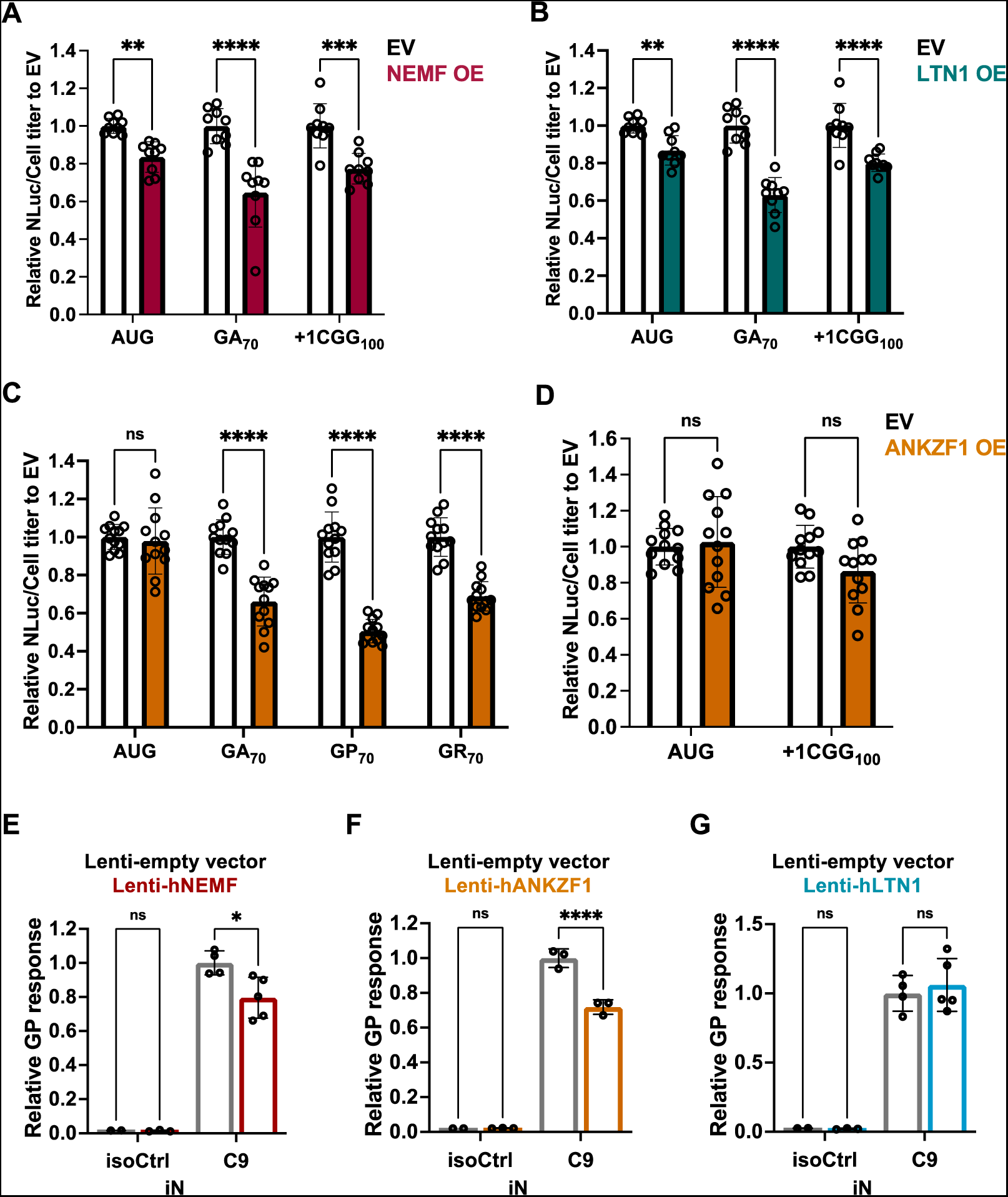
Overexpression of NEMF, LTN1, and ANKZF1 decreases RAN translation from G4C2 and CGG repeats. (**A-B**) Relative expression of AUG-driven no repeats, (G4C2)70 repeats in the GA frame, and (CGG)100 repeats in the polyG frame when overexpressing empty vector (EV) versus hNEMF or hLTN1. (**C**) Relative expression of AUG-driven no-repeat control and (G4C2)70 repeats in the GA, GP, and GR frames when overexpressing empty vector (EV) versus hANKZF1. (**D**) Relative expression of AUG-driven no-repeat control and (CGG)100 repeats in the polyG frame when overexpressing empty vector (EV) versus hANKZF1. Data represent means with error bars ±SD of n = 9-12, ns = not significant; \*\**P* ≤ 0.01; \*\*\**P* ≤ 0.001; \*\*\*\**P* ≤ 0.0001 by two-way ANOVA with Sidak’s multiple comparison test. (**E-G**) Relative GP response of C9 and isogenic control iN treated with lenti-empty vector or lenti-hNEMF, hLTN1, and hANKZF1. The level of GP was measured by MSD. Data represent means with error bars ±SD of n = 3-5, ns = not significant; \**P* < 0.05; \*\*\*\**P* ≤ 0.0001 by two-way ANOVA with Sidak’s multiple comparison test.

## DISCUSSION

Transcribed GC-rich short tandem repeat expansions in *C9ORF72* and *FMR1* form strong RNA secondary structures as either RNA hairpins or G-quadruplexes (38, 39, 42, 43, 120). These structures are critical for repeat-associated non-AUG (RAN) translation (8, 20, 44, 45, 121–123). mRNA secondary structures can impede elongating ribosomes leading to stalling or collision, and activation of? ribosome-associated quality-control pathways (56, 63, 65, 77). To understand how these pathways interplay with RAN translation across GC-rich repeat RNAs, we performed a targeted screen on factors from the mRNA and protein surveillance pathways associated with ribosomal stalling. We identified NEMF, LTN1, and ANKZF1 from the RQC pathway as robust inhibitors of RAN translation on G4C2 mRNA repeats in the GA frame and CGG repeats in the polyG frame. Depletion of NEMF, LTN1, and ANKZF1 increased the abundance of RAN products while overexpression decreased detectable RAN products, with similar genetic compensation effects at endogenous loci in human C9 patient-derived neurons (iNs). Intriguingly, these same factors are upregulated in C9 iNs, suggesting that these pathways may be activated by the translation of expanded repeats. With an N-terminal tagging system, we observed that both G4C2 and CGG repeat sequences generate partially made truncated products, suggesting peptide release associated with RQC degradation pathways activated during repeat translation. Importantly, these effects are mediated not by alterations in the initiation rates (as they persist even when initiation is driven with an AUG initiation codon) or by the charge of the amino acids associated with the peptides (both GA and polyG are uncharged), but by the mRNA structures themselves, as we can abrogate this genetic interaction by changing the repeat sequence to diminish the predicted RNA hairpin structures formed by the CGG repeat element. These data suggest a role for RQC pathways in both RAN translation and AUG-initiated translation of GC-rich structured repeat RNAs, with implications for disease pathogenesis and therapeutic development in this currently untreatable class of disorders.

Several studies implicate RQC activation as associated with neuronal death and human neurodegenerative diseases(88, 124–128). NEMF mutations in mice are sufficient to trigger neurodegenerative phenotypes and variants of NEMF in humans are associated with neuromuscular disease (129, 130). LTN1 mutations in mice also trigger movement disorder phenotypes and motor neuron degeneration (131). Factors from protein surveillance pathways directly associate with ALS/FTD-causing toxic arginine-rich dipeptide repeat (DPR) proteins, GR, and PR (98, 100, 101, 110). AUG-initiated arginine-rich proteins from GR and PR encoding RNAs induce ribosomal stalling in an RNA-independent and DPR protein length-dependent manner in both mammalian cells and GR and PR DPR protein-expressing flies (97, 132). In these cases, ZNF598, NEMF, and LTN1 regulate the expression of GR protein in a fashion that does not require the G4C2 repeat RNA sequence or structure, and these effects are thought to be due to interactions of the charged DPR protein due to the charged arginine residue. Importantly, in these studies, AUG-initiated translation of a GA or GP sequence from a non-repetitive sequence was insufficient to elicit ribosomal stalling. In contrast, here, we observe evidence for ribosomal stalling impacting the expression of GA, GP, and GR generated through RAN translation on G4C2 repeat transcripts that include the native upstream endogenous intronic sequence of *C9ORF72*. We also demonstrate that polyG RAN translation from CGG repeats but not AUG-initiated translation of polyG from a non-repetitive sequence is highly sensitive to RQC factor expression manipulation. As such, our findings suggest that RNA secondary structure formed by GC-rich STRs contributes to ribosomal stalling and activation of downstream surveillance mechanisms. As RAN translation can proceed in multiple reading frames on the same repeat simultaneously, our data cannot rule out a second contribution to ribosomal stalling by rare polyGR translation events on G4C2 or polyR CGG repeats as a stall trigger in other reading frames.

Prior studies utilized P2A-based polycistronic reporter systems encoding two fluorescent proteins to assess translation elongation stalling (91). In these assays, the efficiency of the translation is measured by the ratio of two fluorescent proteins by flow cytometry. However, on GC-rich repeats, the relative loss of expression from a C-terminal tagged fluorescent protein could also result from translational frameshifting, which occurs on both CGG and G4C2 repeats (51, 133). In addition, fluorescent protein systems cannot exclude a contribution to C-terminal tag fluorescent protein expression from internal ribosome entry site (IRES) initiation within these GC-rich repeats as is proposed as a potential mechanism for RAN translation at both CGG and G4C2 repeats (45, 122, 123). To overcome these limitations, we used an N-terminal AUG-V5 tag above G4C2 and CGG repeats to successfully detect short, truncated V5-tagged peptides, which are likely to be released from stalled ribosomes and are resistant to degradation. We also observed that the accumulation of these short peptides is modulated by genetic manipulation of RQC factor expression. Future studies will be needed to see if such peptides are independently toxic and contribute to disease.

Among the modifiers studied, loss of ANKZF1 (yeast Vms1) had the greatest impact on RAN translation product accumulation from both CGG and G4C2 repeats. This factor was not previously assessed with PR or GR-elicited ribosome stalling, presumably because it was thought to act at a very late stage in the RQC pathway. Overexpression of ANKZF1 in mammalian cells selectively decreased expression of GA, GP, and GR from G4C2 repeats and we see a similar of this effect in C9 patient iPSC-derived iNeurons. ANKZF1 is conserved from yeast to humans, where it is thought to act as a hydrolase and/or nuclease to release the nascent peptide chain from the peptidyl-tRNA on the 60S ribosome (95, 96, 134–137). In addition, Vms1 in yeast may have additional roles related to ribosomal stalling that are independent of the canonical RQC pathway, as it can cleave peptidyl-tRNA chains on the leading stalled ribosome independent of 40S and 60S subunit splitting (96). Consistent with this earlier role in stall resolution, Vms1 is found across the entire gradient in yeast polysome profiles, suggesting an association with polysomes as well as isolated and split 60S subunits (96). In the absence of ANKZF1, as with other RQC factors, 60S subunits cannot be recycled for further rounds of translation, leading to a decrease in global translation **(Figure S5)**.

In this work, we highlight NEMF, LTN1, and ANKZF1 as regulators of RAN translation product accumulation from G4C2 and CGG repeats. By detecting truncated products generated from GC-rich transcripts, we suggest that translational stalling occurs within the short tandem repeat region of these mRNAs. We propose that these stalled ribosomes are triggered by the mRNA repeat secondary structure and that they recruit RQC complexes to assist with ribosomal recycling and clearance of aberrant mRNA and proteins (63). How exactly this ribosomal recycling and clearance would work on repeats, however, is somewhat unclear. Classically, the RQC pathway degrades nascent peptide chains after ribosomal splitting and disassembly through a process that is CAT-tailing and NEMF-dependent (85, 86). This CAT-tailing is thought to allow for lysine residues to exit the ribosome and then be (84, 85) ubiquitinated in an LTN1-dependent fashion to allow for targeting of nascent peptide chain to proteasomal degradation (58, 61, 89, 91, 134). However, there is no lysine available for ubiquitination in either of the sequences that serve as RAN translation templates from C9orf72-associated GA, GP, and GR reading frames from the sense transcript or the upstream endogenous 5’UTR, including the CGG repeat, of the FMR1 sequence. Without available lysine, we hypothesize that even when RQC pathways are triggered during RAN translation, the truncated products generated by RQC processing will likely be resistant to ubiquitination and degradation. What roles such truncated degradation-resistant products might play in repeat-associated toxicity will be an important future research direction.

Carboxyl-terminal reporters are often used to measure the efficiency of RAN translation initiation in different settings (44, 45, 49). These approaches face a limitation in that they assume translational elongation is held constant across reporters. As RAN translation potentially happens in multiple reading frames of the same transcript with different rates for both initiation and elongation, interpretation of results from the use of such carboxyl terminal tags as their sole readouts will need to be re-evaluated. Here, we observed largely similar results from both reporters using such a C-terminal tag reporter system as well as from MSD-based assays that directly measure DPR product generation (and as such are not reliant on translation elongation extending completely through the repeat to the NLuc reporter). The increase in generation of both partially made products from stalled ribosomes and complete protein products which contain the C-terminal reporter in the absence of key RQC factors (**Figure 4**) suggests that ribosomal stalling triggers one of two possible events. First, the surveillance pathway might still be triggered, leading to removal of stalled or collided ribosomes from the repeat mRNA but a failure to clear the nascent peptides from the 60S subunit. This would result in accumulation of those stall products and their detection on a denaturing gel. If the repeat mRNA is not degraded during this process, then removal of stalled ribosomes would also allow trailing ribosomes to complete translation through the entire repeat at increased rates, resulting in greater C-terminal reporter signal. Alternatively, the lack of RQC factors may create a longer time window during which stalled ribosomes can resume elongation through the repeat without engagement of the RQC machinery. As such, more ribosomes would make it to the C-terminal reporter and generate complete products.

In summary, we find that RQC pathways inhibit accumulation of RAN translation products generated from GC-rich transcripts. Depletion of NEMF, LTN1, and ANKZF1 increases the accumulation of proteins made via RAN translation from GC-rich transcripts while overexpression of NEMF, LTN1, and ANKZF1 decreases the production of these proteins. These data suggest that augmenting RQC activity could have therapeutic benefits in GC-rich nucleotide repeat expansion diseases and that translational elongation needs to be considered in studies of RAN translation and other initiation-dependent processes.

## DATA AVAILABILITY

All data points are included in the main figures or supplementary data. Raw data used for figure generation are available upon request.

## AUTHOR CONTRIBUTIONS

Y-J.T., I.M., R. S., and P.K.T. conceived the project. Y-J.T. and I.M. designed the experiments. Y-J.T. performed, optimized, and visualized experiments with help from A.K. on plasmid construct and validation; K.J-W and L.P. on GP MSD assay; E.M.H.T. and S.J.B. on patient-derived iPSC iN. I.M. performed Drosophila experiments and assisted with the design and analysis of these studies. X.D. performed polysome profiling experiments. N.B.G. performed initial studies. Y-J.T. wrote the draft of the manuscript with critics from I.M. along with P.K.T., and all authors reviewed and edited the manuscript.

## ACKNOWLEDGMENTS

We thank current and former members of the Todd lab and the Barmada lab for critical discussions, and technical advice in the pursuit of this study. We thank the University of Michigan Vector Core for packaging lentivirus for this study. We thank Dr. Claudio Joazeiro for sharing cell lines and materials for the initial studies and Dr. James Bardwell for his generous support. We thank the University of Michigan Mass Spectrometry-Based Proteomics Resource Facility, especially Dr. Venkatesha Basrur, for the assistance and use of equipment.

## FUNDING

This work was funded by grants from the NIH to P.K.T. (P50HD104463, R01NS099280, and R01NS086810), S.J.B. (R01 NS097542, R01 NS113943, and R56 NS128110) and L.P. (RF1AG062171, R35NS097273, P01NS084974, AG062077) . P.K.T. and A.K. were also supported by the VA (BLRD BX004842). Y-J.T. was supported by the Cellular and Molecular Biology Graduate Program, University of Michigan. LP and KJ-W were supported b by IM was supported by an Alzheimer’s Association Research Fellowship.

## CONFLICT OF INTEREST

The authors declare no competing interests.

---

**Figure S1.**
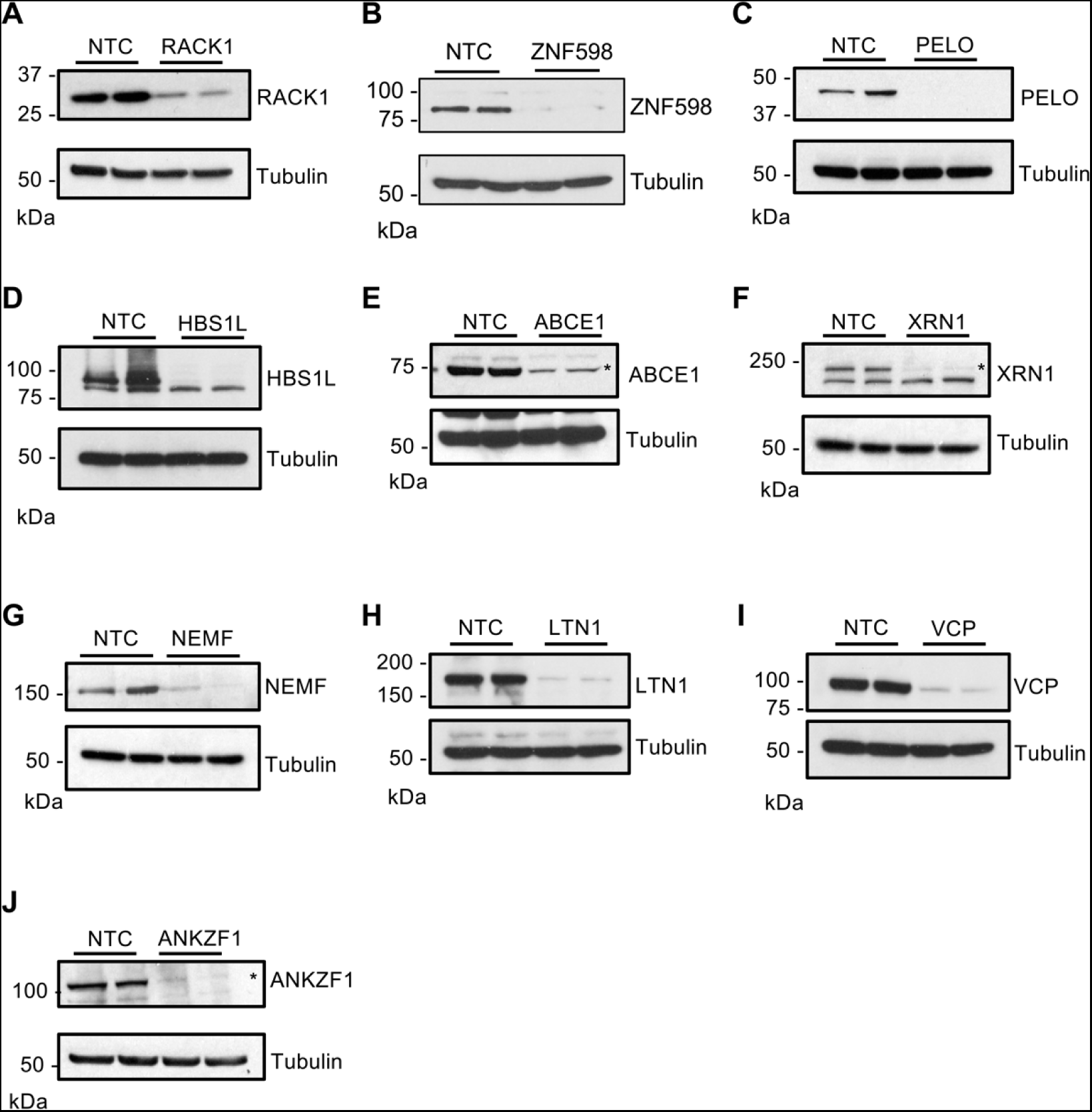
Validation of siRNAs used in targeted screening. (**A-J**) 1nM siRNA transfected in HEK293 of each non-targeting (NTC) control and knockdown of ZNF598, RACK1, PELO, HBS1L, ABCE1, XRN1, NEMF, LTN1, VCP, and ANKZF1

**Figure S2.**
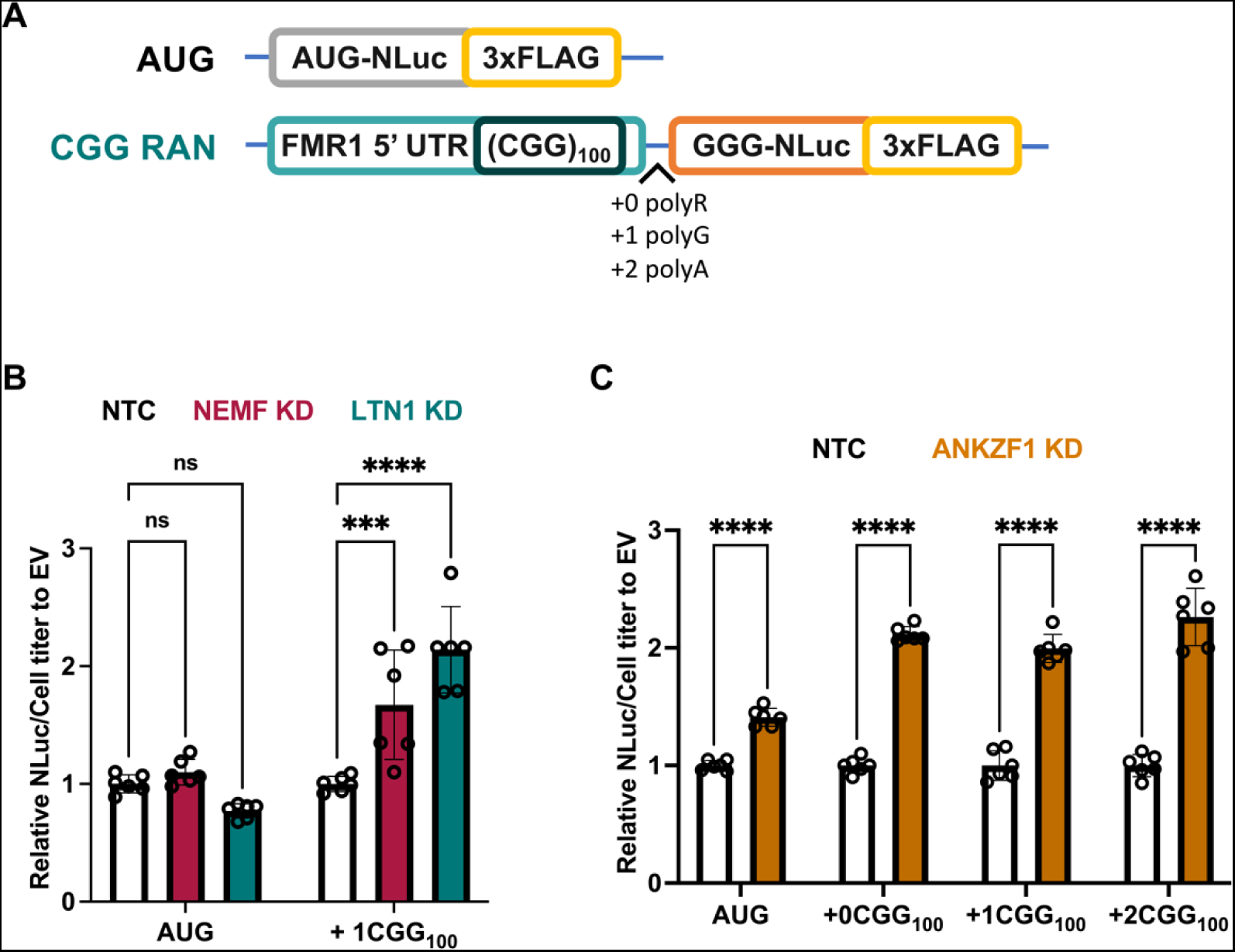
Depletion of NEMF, LTN1, and ANKZF1 enhances CGG RAN translation. (**A**) Schematic AUG-driven and CGG RAN at different reading frame reporters. (**B-C**) Luciferase assays of RAN translation after NEMF, LTN1, or ANKZF1 depletion. All graphs show mean with error bars ± SD. Each N is shown as an open circle (n=6/group). Asterisks above each bar are comparisons of expression between NTC and gene(s) knockdown. ns = not significant; \*\*\**P* ≤ 0.001; \*\*\*\**P* ≤ 0.0001, as determined with one-way ANOVA with Sidak’s multiple comparison test.

**Figure S3.**
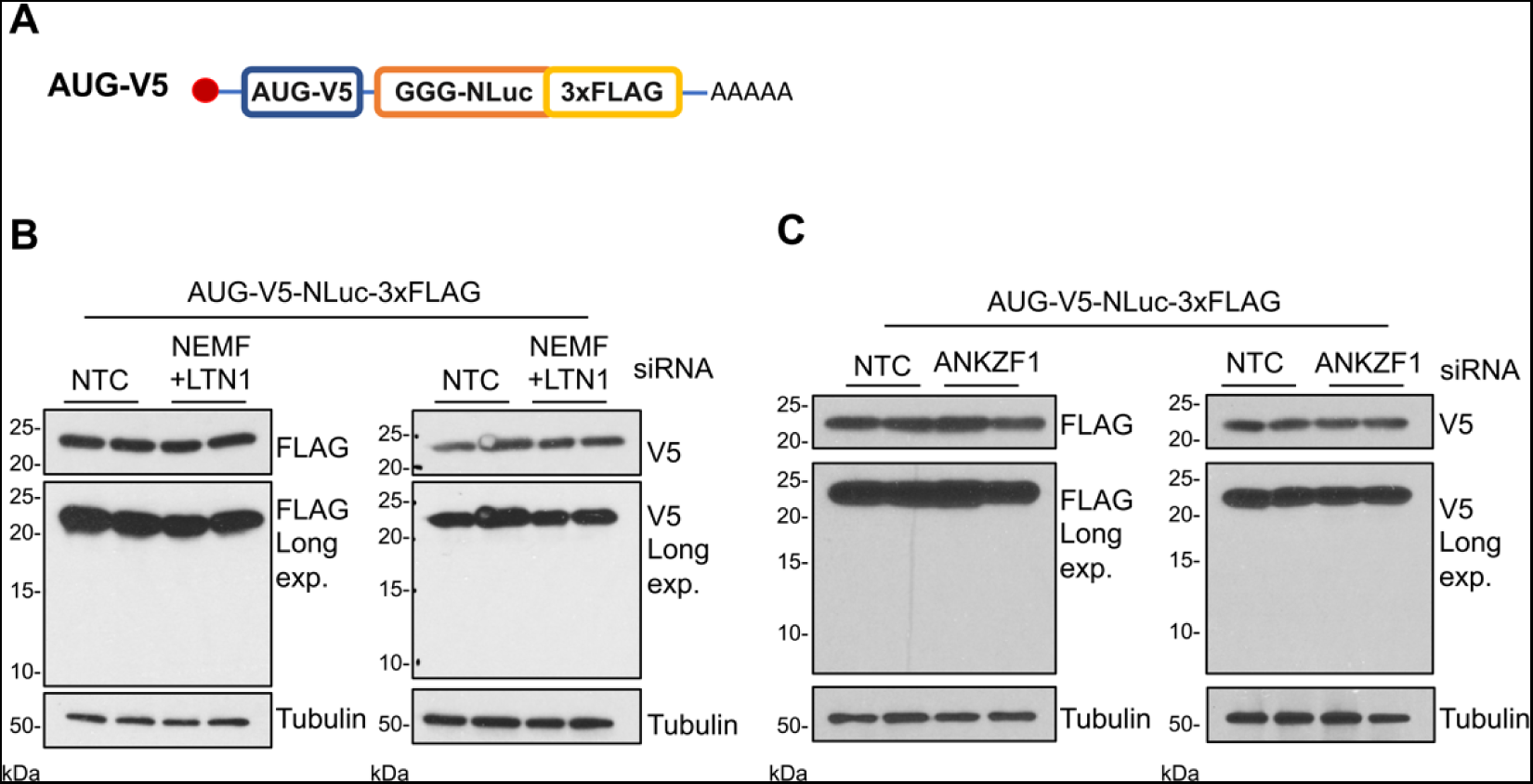
No detection of stall products from constructs lacking GC repeats. (**A**)Schematic of AUG-V5-NLuc-3xFLAG. (**B-C**) Knockdown of NEMF + LTN1, and ANKZF1 in HEK293 with AUG-V5-NLuc-3xFLAG RNA transfection.

**Figure S4.**
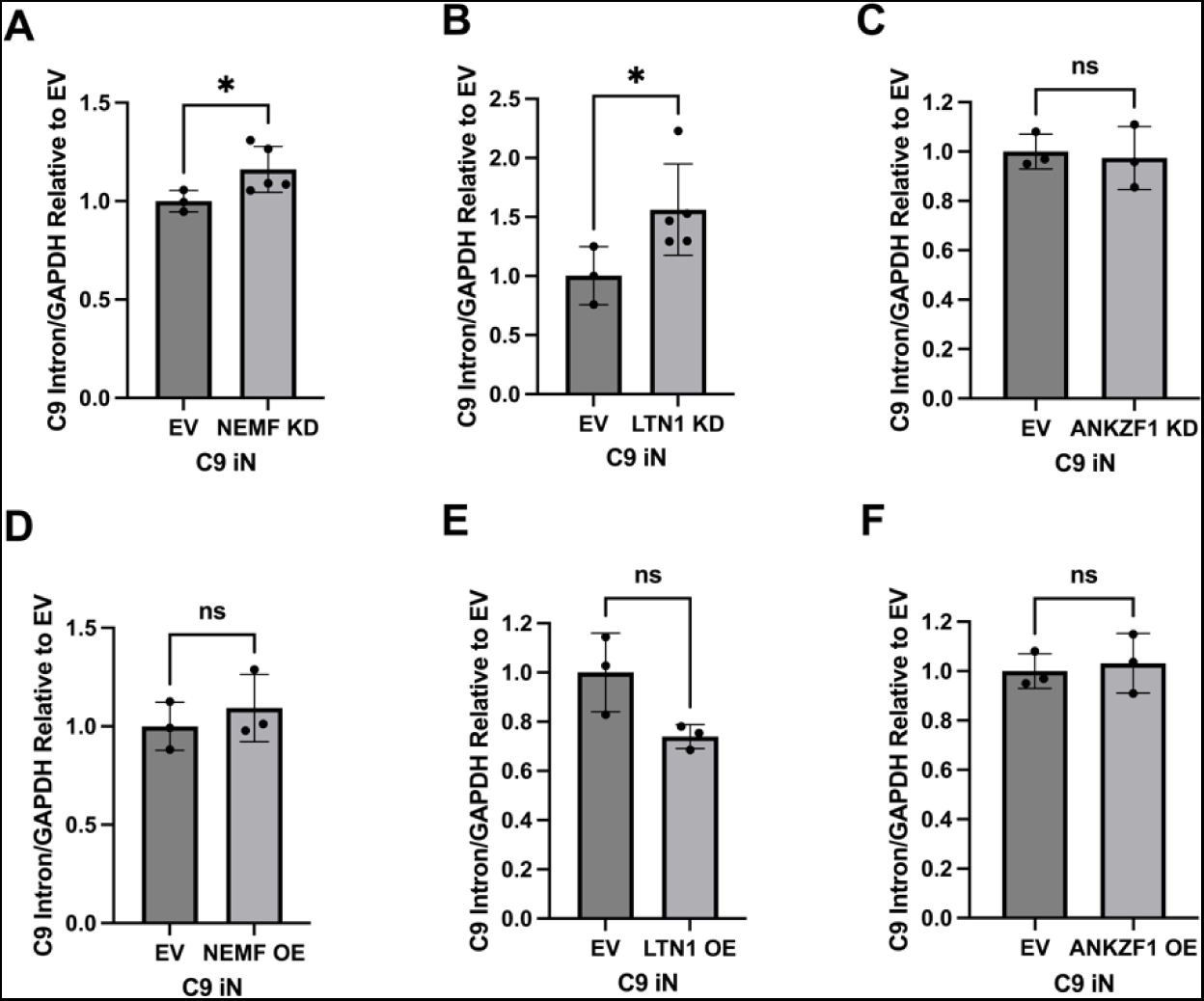
Effect of NEMF, LTN1, and ANKZF1 knockdown and overexpression on C9 transcripts. (**A-F**) Quantification of RNA abundance targeting the C9 intronic region normalized to GAPDH. C9 iN was treated with lentiviruses of empty vector, NEMF, LTN1, or ANKZF1 knockdown or overexpression. Leftover lysates from GP MSD were collected for RNA extraction and qRT-PCR analysis. Data represent means with error bars ±SD of n = 3-5, ns = not significant; \**P* ≤ 0.05 by Student’s t-test.

**Figure S5.**
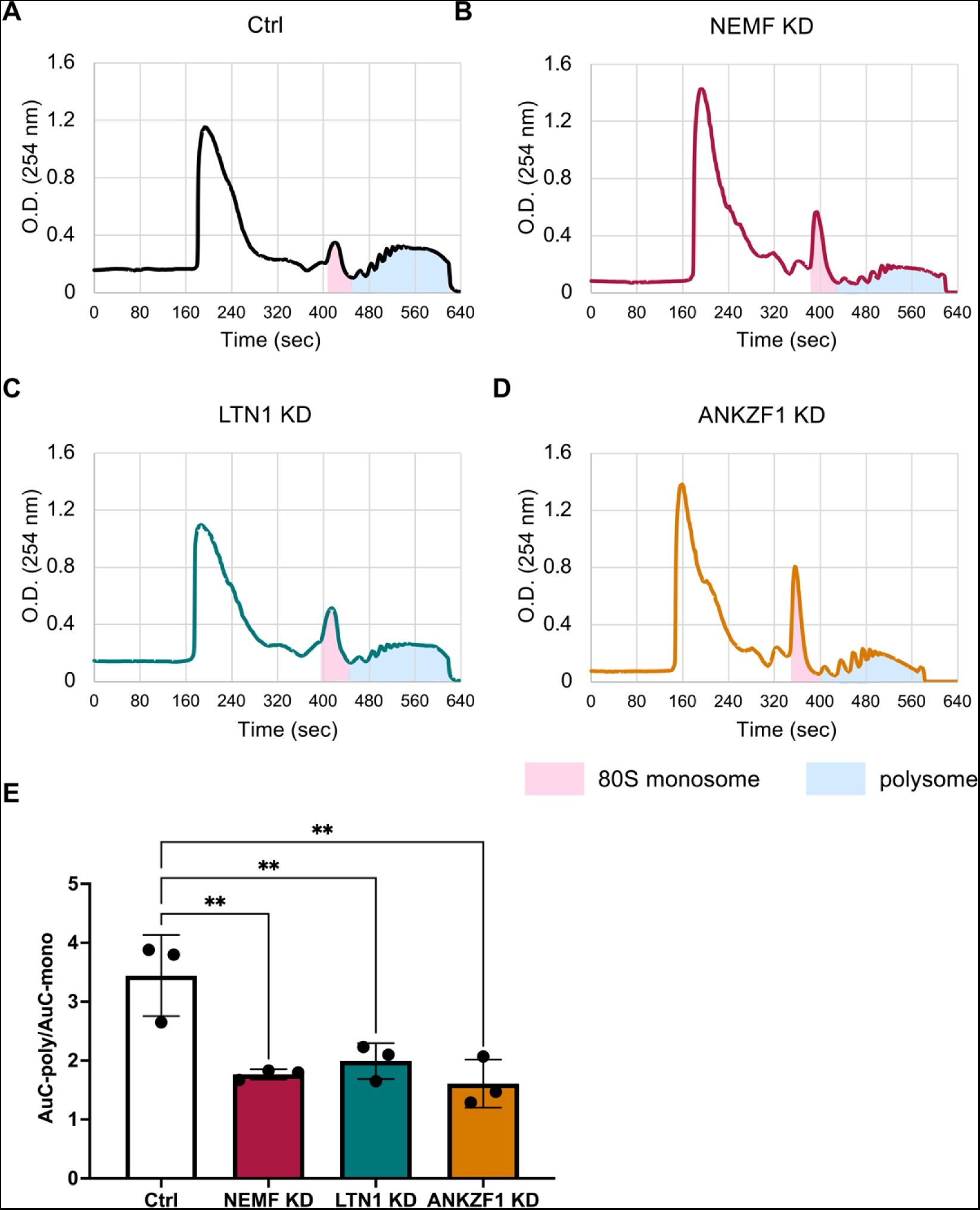
Knockdown of NEMF, LTN1, and ANKZF1 inhibit global translation. (**A-D**) Representative polysome-fractionation profiles of HEK293 lysates transduced with lentivirus of Ctrl KD, NEMF KD, LTN1 KD, or ANKZF1 KD. (**E**) The areas-under-the-curve (AuC) for monosomes and polysomes are shaded pink and blue, respectively. Global translation activity is calculated by normalizing AuC-polysome/AuC-monosome. Data represent means with error bars ±SD of n = 3, \*\**P* ≤ 0.001 as determined with one-way ANOVA with Sidak’s multiple comparison test.

## Notes

### Competing Interest Statement

The authors have declared no competing interest.

